# Sequence and structure of protein binding sites in RNA impact biomolecular condensates

**DOI:** 10.64898/2026.02.24.707737

**Authors:** Sierra J. Cole, Scott R. Allen, Bryan B. Guzmán, Yue Hu, Benjamin M. Stormo, Christine A. Roden, Joanne Ekena, Vita Zhang, Grace A. McLaughlin, Alex W. Crocker, Alain Laederach, Daniel Dominguez, Amy S. Gladfelter

**Affiliations:** Duke University; University of North Carolina; UNIV. DE MONTREAL; University of Minnesota; University of North Carolina at Chapel Hill

## Abstract

Biomolecular condensates are central to subcellular compartmentalization. Although many condensates contain and regulate RNA, research has primarily focused on protein interactions. Here, we investigate RNA–protein interactions underlying cell cycle–regulating condensates in the multinucleate fungus *Ashbya gossypii*. These condensates form through interactions between G1 cyclin mRNA *CLN3* and RNA-binding protein Whi3, which was predicted by homology to recognize a five-nucleotide motif repeated within the transcript. Natural variation in motif number in *Ashbya* strains led us to hypothesize that binding site valence may influence condensate properties. Using unbiased binding assays, we determined the preference of *Ashbya* Whi3 protein for specific primary RNA sequence and mutated individual Whi3-binding sites within *CLN3* mRNA. Mutants exhibited distinct condensate properties despite having the same valence in terms of binding site number. Mutations altered the saturation concentration (Csat) and dense phase concentration of RNA and protein in cell-free reconstitution experiments. A subset of mutants, showed reduced number of condensates and deregulation of the cell cycle in cells. We also find that enhanced availability of single-stranded RNA can compensate for loss of binding sites. Together, these data indicate that differences in RNA protein binding-site context and not simply valence plays a critical role in determining condensate properties.

## Introduction

The physical organization of the cytosol has been a source of fascination since the19th century (Hyman & Simons, 2012; Wilson, 1899). Advancements in multiple fields established that biomolecules such as RNA, DNA, and proteins can undergo phase separation into membraneless liquid-like assemblies known as biomolecular condensates (Banani et al., 2017; Brangwynne et al., 2009; Li et al., 2012). A hallmark of molecules that form condensates is the ability to form multiple reversible interactions with other molecules, known as multivalency (Banani et al., 2017; Li et al., 2012). With regard to proteins, these interactions include an array of non-covalent associations such as electrostatic, hydrophobic, dipole-dipole, cation-π, or π-π interactions (Brangwynne et al., 2015; Holehouse & Alberti, 2025; Perry, 2019). Proteins that phase separate often contain intrinsically disordered regions (IDRs), which lack a single structure, allowing multiple different conformations, thereby helping to increase potential valency (Borcherds et al., 2021). Significantly less is understood about the properties of RNA that dictate which molecules will drive or enrich condensates. Like IDRs, RNAs can convert between multiple different structural states (Bonilla et al., 2024; Nakano & Sugimoto, 2016). The negative charge of RNA can promote non-specific interactions with positively charged proteins to form condensates; however, there are also many cases in which specific protein and RNA interactions are essential for the formation of condensates (Elguindy & Mendell, 2021; Iserman et al., 2020; Nordenskiöld et al., 2024; Roden & Gladfelter, 2021; Zhang et al., 2015). In this paper, we interrogate the specific interactions between protein and RNA that give rise to biomolecular condensates using a well characterized and biologically-relevant RNA and protein condensate from *Ashbya gossypii*.

*Ashbya* is a multinucleate filamentous fungus that exists in the digestive systems of insects across North America (Ashby & Nowell, 1926; Wendland & Walther, 2005). Nuclei in these cells undergo mitosis asynchronously despite residing in a shared cytoplasm, indicating spatial restriction of cell cycle factors to keep nuclei cycling independently of one another (Gladfelter, 2006). A key mechanism of asynchrony lies in Whi3, an RNA-binding protein. Whi3 binds and forms condensates with the G1 cyclin RNA, *CLN3*, controlling *CLN3* localization and expression (Geisterfer et al., 2024; Lee et al., 2013). Reducing Whi3’s ability to bind to itself or to *CLN3* greatly decreases condensate formation while increasing synchrony, demonstrating the importance of RNA-protein binding and condensate formation for local cell cycle control (Lee et al., 2013). Natural variation in *Ashbya* strains isolated from different climates show sequence changes in both Whi3 and *CLN3* that promote distinct condensates at specific temperatures (Stormo et al., 2024). Variability in the number of predicted Whi3 binding sites in *CLN3* led us to examine how protein binding site number and context in *CLN3* may contribute to RNA-protein interactions and the properties of condensates. We show that despite these motifs having identical sequences, the sites are not functionally equivalent for molecular interactions and condensate formation.

## Results

### Whi3 protein has preferential binding to specific primary sequence motifs

Previous work has shown that the *S. cerevisiae* Whi3 homolog binds to a UGCAU motif and this sequence appears conserved in known Whi3 targets in *Ashbya*; however, this sequence can arise by chance and it had not been explicitly tested if *Ashbya* Whi3 indeed binds this sequence or if there are other interaction motifs that contribute to Whi3-*CLN3* condensates (Lee et al., 2013; Riordan et al., 2011). We used RNA bind-n-seq in which varying concentrations of His-tagged Whi3 were mixed with 20 nt randomized RNA oligonucleotides (Lambert et al., 2014). Whi3 was then pulled down and the oligos bound were recovered, sequenced, and analyzed for 5-mer or 6-mer motif enrichment (R value) which is calculated by dividing the number of reads in the presence of protein by a input control with no protein (Dominguez et al., 2018; Lambert et al., 2014). When considering 5-mer motifs, we found that Whi3 bound most strongly to UGCAU as predicted by homology to *S. cerevisiae* at 250 nM Whi3; however, there were other nucleotide motifs with significant R values at 250 nM Whi3 and reducing Whi3 concentration results in less specificity for UGCAU (Sup Figure 1A-C). Expanding to 6-mers, we see that the nucleotides surrounding the canonical UGCAU motif greatly affect the ability of Whi3 to bind, as UUGCAU has an R value of 14.61. In contrast, GUGCAU exhibits an R value of only 4.03, which is lower than some 6-mers that do not contain the UGCAU motif (Figure 1A). We wondered if this difference was purely a primary sequence preference or if these different 6-mers may vary in predicted structure, affecting Whi3 binding since RRMs frequently prefer unstructured RNA (Auweter et al., 2006; Lunde et al., 2007). Using the top 500 enriched 6-mers, we a slight preference for unstructured 6-mers though primary sequence appears to play a more significant role (Figure 1B). To maintain consistency with previous papers, we will refer to UGCAU as the preferred Whi3 binding site sequence throughout the rest of this manuscript. *CLN3* mRNA contains five of these binding sites, four of which are in the 5’ UTR and one in the coding sequence.

**Figure 1.**
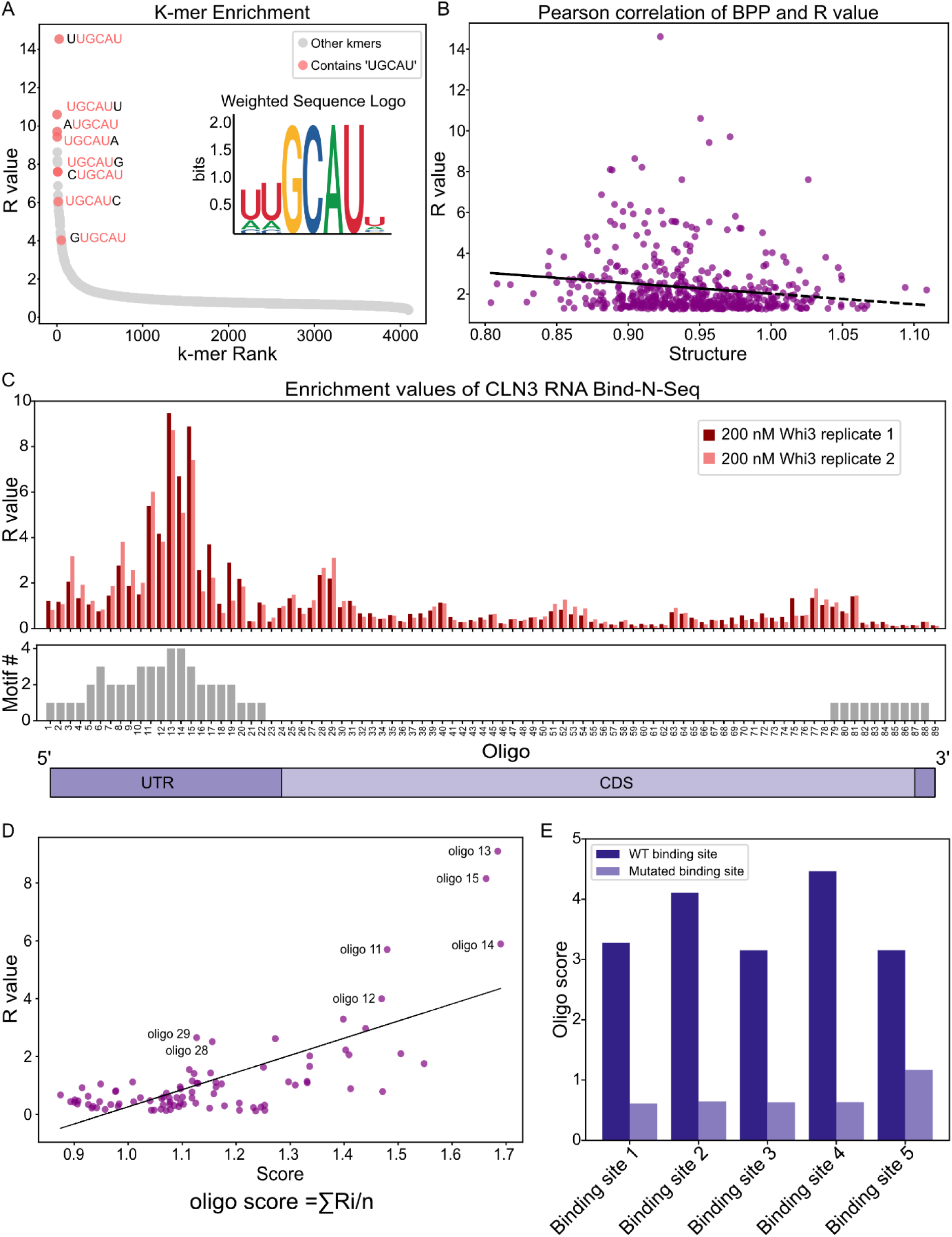
Determining primary RNA interaction sites for Whi3. A) A graph showing which 6 nucleotide motifs were most enriched in the random oligo pool RNA bind-n-seq experiment using 250 nM Whi3. UGCAU containing motifs are highlighted in red. This experiment was done with two biological replicates. B) Pearson correlation between base-pairing probability of 6-mers in random oligo pool RNA bind-n-seq and R value score. Pearson’s r = -0.16 p-value = 0.000274 C) Visualization of R values for *CLN3* oligo-specific RNA bind-n-seq experiment. The graph below shows the number of UGCAU sites marked for each oligo. A visual representation of the different regions of the transcript is shown below this. D) Pearson correlation of *CLN3* oligos comparing the oligo score obtained from averaging the R value score of 6-mers in each oligo and the R value score of *CLN3* oligo-specific RBNS. Pearson’s r = 0.72 p-value = 1.48X10^-15^ E) Comparison of the oligo score of the WT binding site within the *CLN3* sequence (+/- 5 nucleotides on either side) and the mutations meant to eliminate Whi3 binding.

To confirm that Whi3 is primarily binding to the UGCAU sites within *CLN3* mRNA, we ran another RNA bind-n-seq experiment using 200 nt oligos tiled across *CLN3*. As expected, there was a significant peak in R value in Whi3 binding to oligos that contained the UGCAU sites, which was further enhanced if the oligo contained multiple binding sites (Figure 1C). To better understand why certain oligos were enriched in the Whi3 pull-down, we scored the tiled *CLN3* oligos using the 6-mer R value scores from the random oligo pool experiment. To do this, we averaged the R value scores of each 6-mer available in the *CLN3* oligo. We found a positive correlation between the oligo score and R value score, providing further evidence that a driver of Whi3 binding is primary sequence (Figure 1D). Interestingly, we also found Oligo 28 and Oligo 29 right at the beginning of the coding sequence of the *CLN3,* which does not contain UGCAU, yet do show modest R values with a similar score to some of the oligos toward the 3’ end, which do contain UGCAU. Oligo 28 and 29 do however contain two AGCAU motifs within 11 nucleotides of each other. This AGCAU motif shows an R value score of 5.59 as opposed to UGCAU, which has an R value score of 8.48. It is possible that even though Whi3 does not have as strong an affinity to these sites, having these two lesser sites close together makes up for the lack of the precise UGCAU. In addition to scoring the oligos via primary sequence, we also looked at each total oligo’s predicted minimum free energy (MFE) and ensemble diversity (ED) using Vienna RNA to assess if structures were associated with binding (Gruber et al., 2008). We did not see a significant correlation between R value and predicted structure, similar to the results we saw with the random oligo pool at 250 nM Whi3 (Sup Figure 1B-D). These results suggest that in the context of the short oligos used in these experiments, Whi3 preferentially binds to UGCAU motifs and that this interaction can be driven by primary sequence.

These combined experiments indicate there are 5 dominant Whi3 binding sites (WBS) on *CLN3*. We next mutated individual binding sites to CUAGU for the first four sites, which are in the 5’ UTR, and to UACAC for the fifth site in the coding sequence (to retain protein coding) to determine if and how the context of individual sites contributes to Whi3’s ability to form condensates. Using the same oligo scoring approach to assess the individual binding sites within *CLN3*, we showed that each of the mutations decreased this score to approximately 0.60 for all binding sites except Whi3 binding site 5 mutant, *wbs5* (Figure 1E), which was constrained in design to preserve the coding sequence for ultimate functional analysis in cells. The mutant sequences were used to generate *in vitro* transcribed RNAs for *in vitro* phase separation assays and integrated into cells for functional analysis.

### Binding site mutants show differences in *in vitro* condensation

We next analyzed whether all Whi3 binding sites contributed equally to promoting condensation of Whi3 *in vitro* since it is known that Whi3 protein condensates require interactions with RNA to form at physiological salt and protein concentrations (Zhang et al., 2015). Using a range of starting concentrations of protein and RNA, we found that both WT *CLN3* and the binding site mutants could form condensates under a variety of different conditions (Figure 2A). Unexpectedly, many of the mutants formed condensates at lower concentrations of protein and RNA (Csat) than WT despite having fewer binding sites and thus reduced valency. *wbs1*, *wbs2*, *wbs4*, and *wbs5* condense at 2 nM RNA and 200 nM Whi3, whereas no condensates are seen at these concentrations for WT RNA (Figure 2A). The *wbs2* mutant could even form condensates at 100 nM Whi3, unlike any other mutant or wild-type sequence (Figure 2A). In contrast, *wbs3* and the no binding site mutant (*nwbs*, previously called 5m mutant in Lee et al 2015 (Lee et al., 2015)) had much higher Csat values for RNA and/or protein (Figure 2A). These results showed that although the mutant alleles, aside from *nwbs*, have the same number of binding sites, they differ in the degree to which they impact phase behavior indicating that it is not the number of binding sites or valence of these sites that is relevant for determining the Csat for this system.

**Figure 2.**
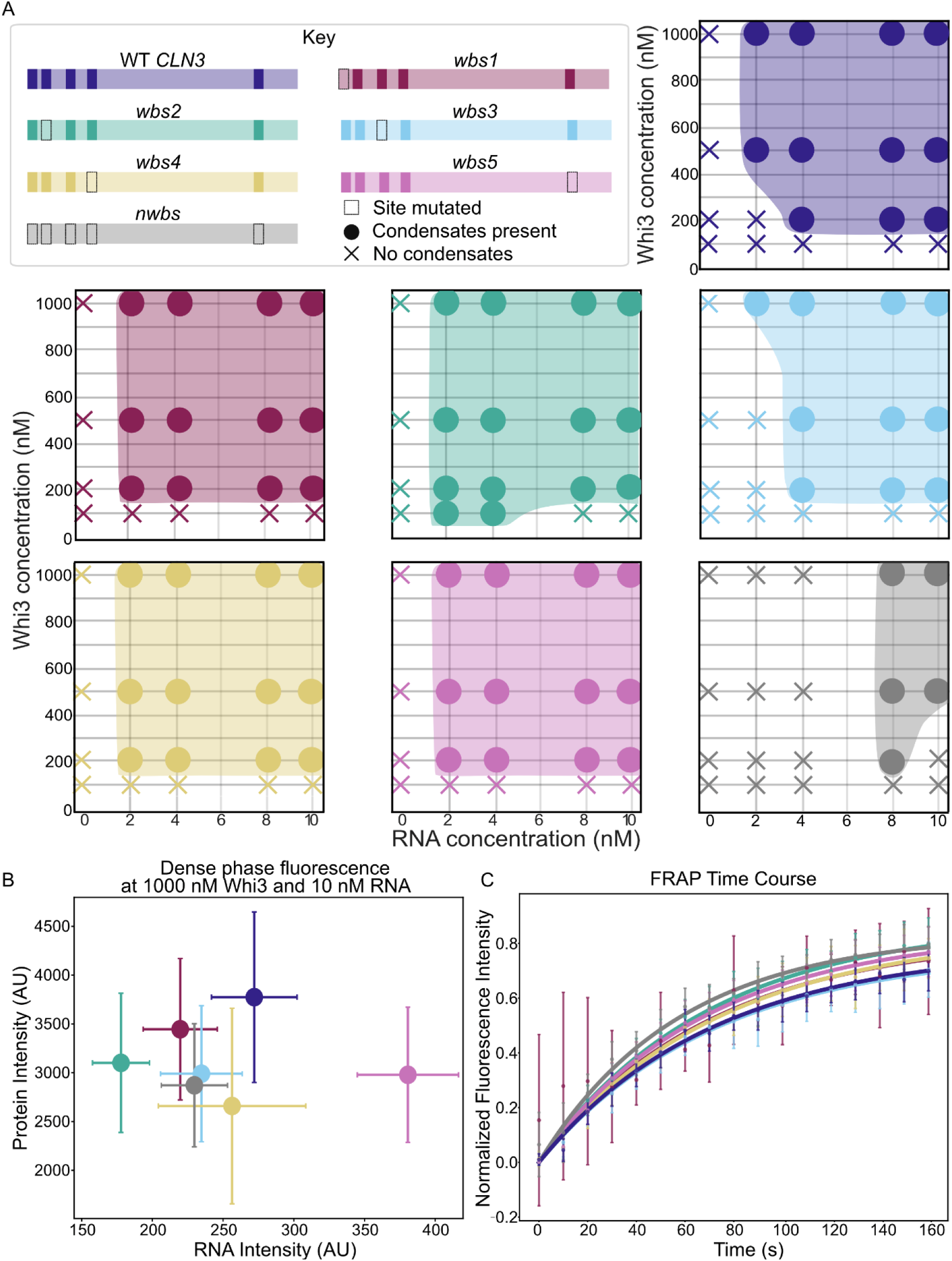
Mutants show distinct differences in condensate properties *in vitro* A) Phase diagrams of WT and mutant *CLN3* showing multiple tested Whi3 and RNA concentrations to determine where condensates form. Conditions where condensates form are represented by circles and Xs represent conditions with no detectable condensates. B) Comparison of relative fluorescent protein intensity and RNA intensity in the dense phase at 1000 nM Whi3 and 10 nM RNA. Error bars show standard deviation n = 600 condensates per RNA type C) FRAP time course with error bars representing standard deviation with 2 biological (independent protein purification and RNA transcription) replicates and 3 technical replicates per biological replicate. Tau values or ½ recovery time was compared using with p-values calculated using Mann-Whitney comparing WT to mutants (*wbs1* p=0.421*, wbs2* p=1.00*, wbs3* p= 0.792*, wbs4* p=1.00*, wbs5* p=0.730*, nwbs* p=0.413).

We proceeded to further characterize differences in the condensates formed by the mutants. We first determined the relative amount of protein and RNA recruited to the dense phase of condensates made with bulk concentrations of 1000 nM Whi3 and 10 nM *CLN3* using fluorescently-labeled components. We found that all mutants showed significantly less Whi3 protein in the dense phase than WT. Most of the mutants also had significantly less RNA present in the dense phase compared to WT, except for *wbs5*, which showed a significantly higher amount of RNA in the dense phase than WT and the mutants (Figure 2B). While all of the condensates appeared spherical in morphology, *wbs1*, *wbs2*, *wbs3*, and *wbs4* had smaller size condensates than WT while *wbs5* and *nwbs* had larger condensates (Sup Figure 2A and B). Previous work in this system had shown that Whi3 and *CLN3* condensates with a higher protein:RNA ratio had faster recovery time of protein during fluorescence recovery after photobleaching (FRAP) experiments and fused more readily while RNA did not show recovery in any of these experiments (Zhang et al., 2015). All of the mutants had significantly different protein:RNA ratio than WT, with *wbs1* and *wbs2* having a higher protein:RNA ratio and the rest of the panel having a lower protein:RNA ratio (p<0.001). Despite differences in the level of recruitment of RNA and protein to the dense phase, there was no significant difference in the recovery time between the mutant RNAs and WT after performing FRAP experiments on labeled Whi3 (p>0.1, Figure 2C). Overall, we found that mutations to individual binding sites affected condensate properties, indicating an unexpected complexity at the scale of protein-RNA interactions in directing phase behavior and condensate properties.

### A subset of binding sites required for condensate formation and function in cells

We next assessed how mutant alleles impacted condensate formation and nuclear cycling *in vivo*. In an *Ashbya* meGFP-tagged Whi3 strain, we replaced the endogenous *CLN3* with either a WT control or one of the binding site mutants. After confirming the genotype of the strains via sequencing, we measured Whi3 foci formation in live cells and found that most cells contained Whi3 puncta, although both *wbs2* and *nwbs* showed significantly less Whi3 in foci compared to WT (Figure 3A and B). To evaluate if these differences are due to different RNA levels, we performed RNA FISH on cells expressing the mutant constructs. We found that while *nwbs* did show somewhat lower expression, *wbs2* did not have lower numbers of transcript, and thus this is not an explanation for the diminished condensates (Figure 3B). Additionally, *wbs1* showed a slight increase in the amount of transcripts present compared to WT yet did not show a significant change in the fraction of Whi3 in puncta. We next assessed how the mutants impacted cell cycle progression and nuclear asynchrony. By comparing the spindle pole bodies of adjacent nuclei, we found that *wbs1*, *wbs2*, *wbs5*, and *nwbs* were all significantly more synchronous than WT (Figure 3E). *wbs4* has fewer nuclei in mitosis and *wbs5* has more nuclei in mitosis but otherwise cell cycle progression was not majorly altered suggesting synchrony is not primarily due to arrests in the cell cycle (Figure 3D). In summary, mutating a single binding site within *CLN3* is sufficient in some cases to alter condensate formation and perturb cell cycle synchrony in cells but again, not all binding sites are equivalent in their impact on Whi3 assembly or function.

**Figure 3.**
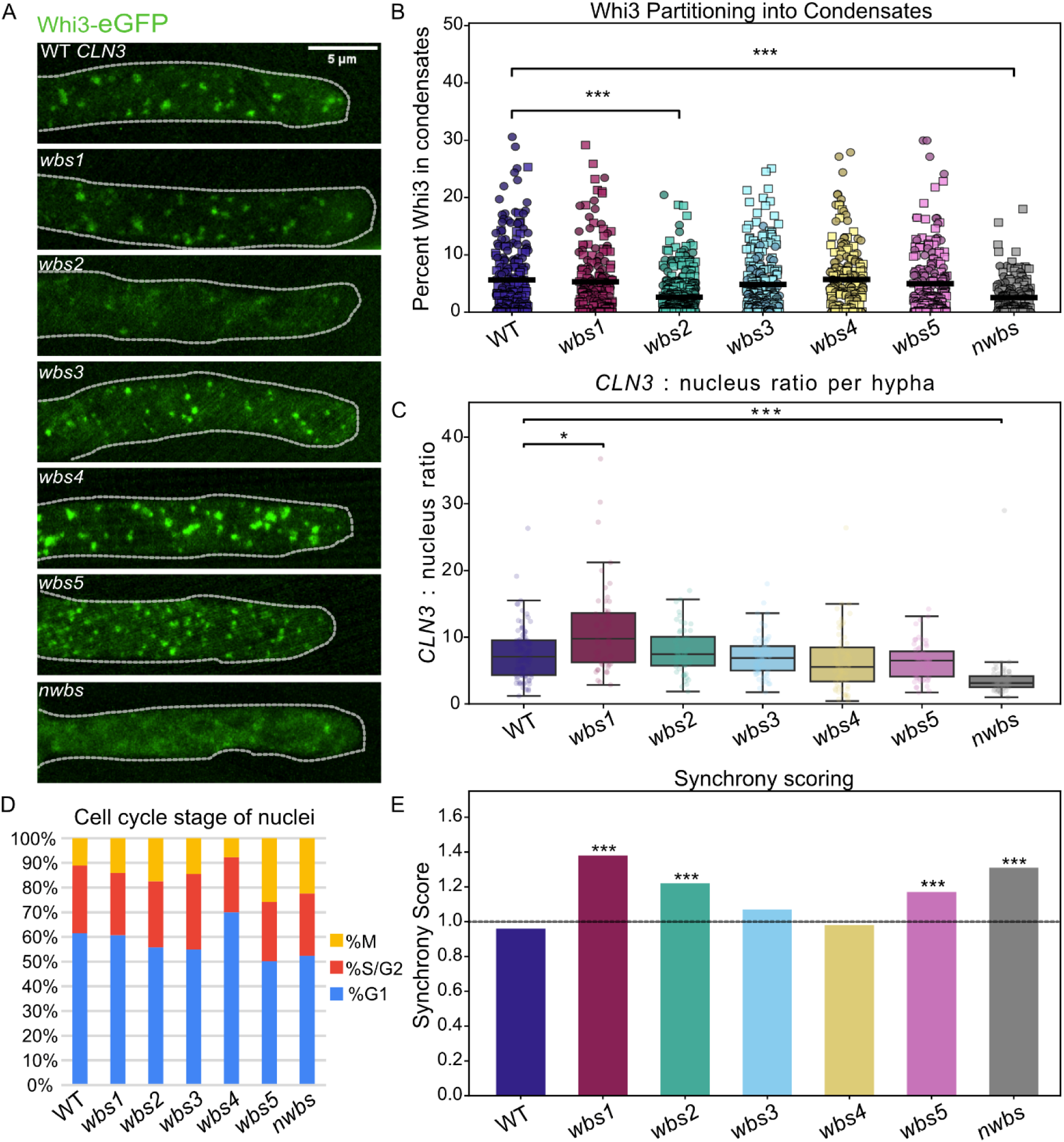
Mutants show distinct differences in condensate properties *in vivo.* A) Representative images of hypha for each mutant. B) Graph showing the quantification of Whi3 signal in foci normalized by the total Whi3 signal in the hypha, with circles and squares indicating different biological replicates. Mean shown via black line and significance calculated using the Kruskal-Wallis test and Dunn Q test *** p < 0.001 n = 100 hyphae per replicate. C) Boxplot showing the quantification of *CLN3* FISH per nucleus in hypha. Mean shown via black line and significance calculated using the Kruskal-Wallis test and Dunn Q test * p < 0.05 and *** p < 0.001. WT n = 91 hyphae, *wbs1* n = 53 hyphae, *wbs2* n = 52 hyphae, *wbs3* n = 67 hyphae, *wbs4* n = 63 hyphae, *wbs5* n = 60 hyphae, and *nwbs* n = 58 hyphae. D) Percent nuclei in different cell cycle stages for different *CLN3* strains. E) Synchrony scoring of different *CLN3* strains *** p < 0.001. WT n = 43 hyphae, *wbs1* n = 60 hyphae, *wbs2* n = 104 hyphae, *wbs3* n = 71 hyphae, *wbs4* n = 79 hyphae, *wbs5* n = 117 hyphae, and *nwbs* n = 88 hyphae

### Mutants do not generate large-scale structural differences

One possibility as to why the mutants behave differently from one another is that altering the primary sequence of the binding sites has disparate ancillary consequences on RNA secondary structure, conceivably even at a distance from the primary site. Vienna RNA predictions showed <10 kcal difference in the predicted MFE between the mutants and WT, except for *nwbs* (Gruber et al., 2008). Additionally, co-fold predictions did not show any major changes in expected dimerization propensity, except *nwbs* (Sup Table 2). To experimentally determine if the RNA secondary structure was significantly changed by the introduced mutations, we performed DMS-MaP (Dimethly Sulfate Mutational Profiling) to compare the WT RNA and the mutants’ secondary structure. We did not observe any significant changes in the DMS reactivity across the mutants, except for *nwbs*, as can be seen by the linear regression of mutants compared to WT in Figure 4A in which we consider R^2^≥0.9 as being not significantly different. However, when we computed the average base pairing probability (BPP) of individual binding sites in WT RNA using two biological replicates, binding site 1 has a higher probability of being double stranded than binding site 2 and 4 (p<0.05, Figure 4B). Like the DMS-MaP data, there was no significant difference between the mutants and the WT RNA when compared by native gel electrophoresis, where mutants showed comparable RNA dimerization capacity (Figure 4C). This indicates that *in vivo* and *in vitro* changes to condensate behavior are not due to large-scale detectable changes in RNA structure caused by the mutations or due to a change in the strength of the RNA-RNA interactions.

**Figure 4.**
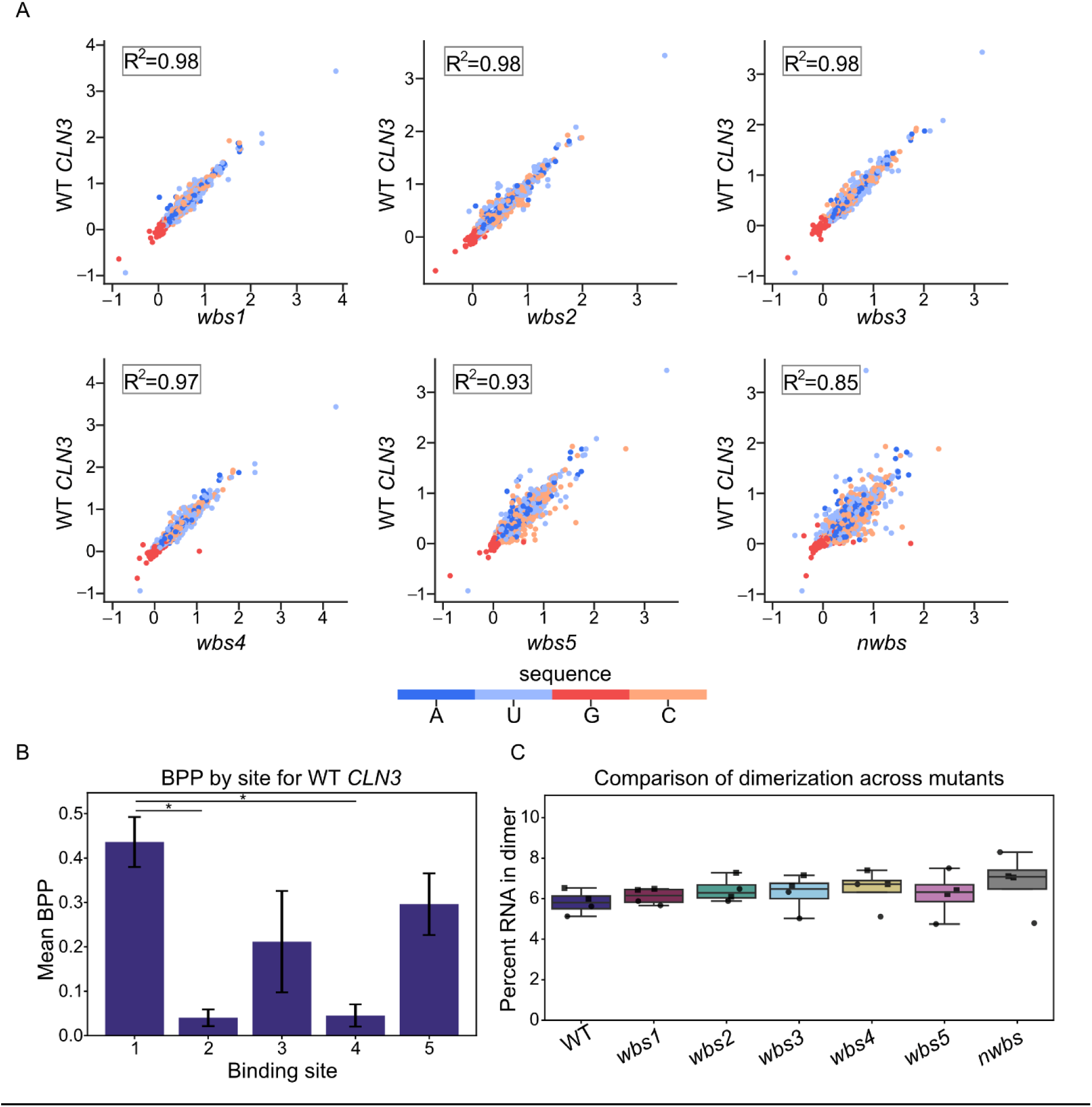
Analysis of secondary structure and dimerization of mutants compared to WT. A) Linear regression comparing normalized DMS reactivity between mutant and WT *CLN3.* The R^2^ values are Pearson’s correlation coefficient. B) Average base-pairing probability (BPP) of the five nucleotides in individual binding sites within WT *CLN3* with error bars representing standard deviation of probability. The difference determined using Welch’s t-test (* p<0.05). C) Quantification of RNA in the dimer band from native RNA gel with squares and circles representing individual biological replicates. There was not significant difference found between strains using one way ANOVA p = 0.838.

### Melting and refolding of mutant RNAs elicits specific effects on condensation

Experiments above showed that mutating individual Whi3 binding sites produces distinct condensate properties *in vitro* and *in vivo*. Although we did not see a change in the global structure of the mutant RNAs compared to WT, we do see a difference in the BPP of binding sites which may influence Whi3-RNA interactions differentially. To further examine possible structure-based differences at play between the sites, we melted the RNA to disrupt co-transcriptionally formed secondary structure prior to mixing with Whi3 and compared condensate properties to those formed with unmelted RNA that folded co-transcriptionally. In parallel, melted RNAs were allowed to refold slowly before being added to protein to assess the effects of re-established or potential distinct structures from those formed co-transcriptionally. Consistent with Figure 2B, all unmelted mutant RNAs recruited significantly less Whi3 to the dense phase than WT (Figure 5A and 5C). However, melting the RNA reduced these differences, with mutants recruiting Whi3 at levels closer to WT when melted. This suggests that overall RNA structure is impacting Whi3’s affinity or accessibility to binding sites since removing co-transcriptionally formed secondary structures from the RNA enhances Whi3 recruitment to condensates. Similarly, melting RNAs increased recruitment of the RNA to the dense phase for *wbs1*, *wbs3*, *wbs4* and *nwbs* compared to unmelted, but not for *wbs5*.

**Figure 5.**
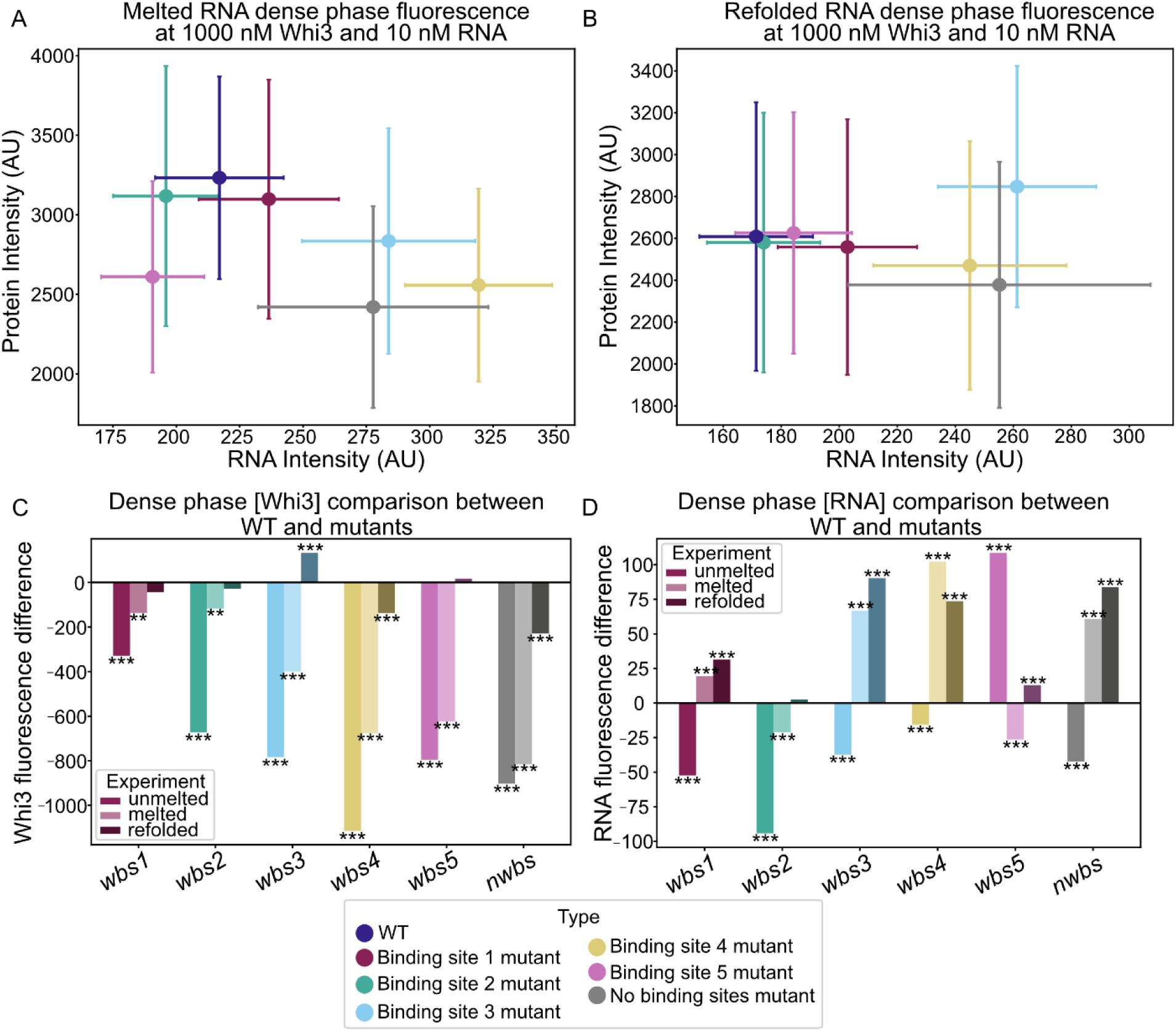
Melting and refolding experiments show that both primary sequence and structure influence condensate properties. A) Comparison of relative fluorescent protein intensity and RNA intensity in the dense phase at 1000 nM Whi3 and 10 nM RNA with RNA that was melted directly before being added to protein. Error bars are showing standard deviation. B) Comparison of relative fluorescent protein intensity and RNA intensity in the dense phase at 1000 nM Whi3 and 10 nM RNA with RNA that was melted and refolded before being added to protein. Error bars are showing standard deviation. N = 600 condensates per RNA type for both melting and refolding experiments. C) Visualization of dense phase fluorescent protein intensity difference between WT and labeled mutants comparing unmelted RNA, melted RNA, and refolded RNA. D) Visualization of dense phase fluorescent RNA intensity difference between WT and labeled mutants comparing unmelted RNA, melted RNA, and refolded RNA. The significance for both the difference in protein and difference in RNA was calculated using the Kruskal-Wallis test and Dunn Q test * p < 0.05, ** p < 0.01, and *** p < 0.001

Interestingly, refolded RNAs did not simply mirror the unmelted phenotype in Figure 2B. Instead, refolded mutant RNAs recruited Whi3 at levels comparable to, or in the case of *wbs3* exceeding WT (Figure 5B and 5C). For RNA recruitment the difference between unmelted and refolded are even more pronounced with all exceeding or matching wild-type levels in the dense phase. These differences likely reflect distinct folding pathways since the unmelted RNAs were folded co-transcriptionally at 37 °C, whereas refolded RNAs were folded post-transcriptionally at 25 °C. Together, these results from unmelted, melted, and refolded RNA indicate that RNA folding and structure play a critical role in determining condensate properties and again very small changes in primary sequence can alter phase behavior depending on the folding history and state of the RNA. In summary, analysis of these mutants indicates that it is not merely the number of Whi3 binding sites that determines mesoscale condensate properties and that individual binding sites can make distinct contributions leading to complex effects depending on their presence or absence.

## Discussion

This study shows how small changes in an RNA sequence in specific protein-RNA interaction sites can influence condensate properties. Unexpectedly, we found RNA transcripts that contained the same number of protein-binding sites but in different locations in the transcript had different phase behaviors from one another which can be seen in Table 1. We determined that none of the mutant RNA sequences showed significant large-scale structural changes or any change in dimerization propensity (Figure 4), leaving the molecular basis for the varied behavior mysterious. Still, the melting and refolding experiments do indicate that overall RNA structure can contribute to condensate properties in ways that are in part related to the sequences. These data show that the impacts of RNA-binding site mutations can have multiscale effects and that the structural properties of RNA can confound predicting the impact of loss or gain of binding sites. The consensus sites for RBPs can be readily gained and lost in evolution, often through synonymous mutation or changes in UTRs, and this study shows that the potential impact of such alterations on condensation of an RBP may extend beyond simple valence effects.

**Table 1:**
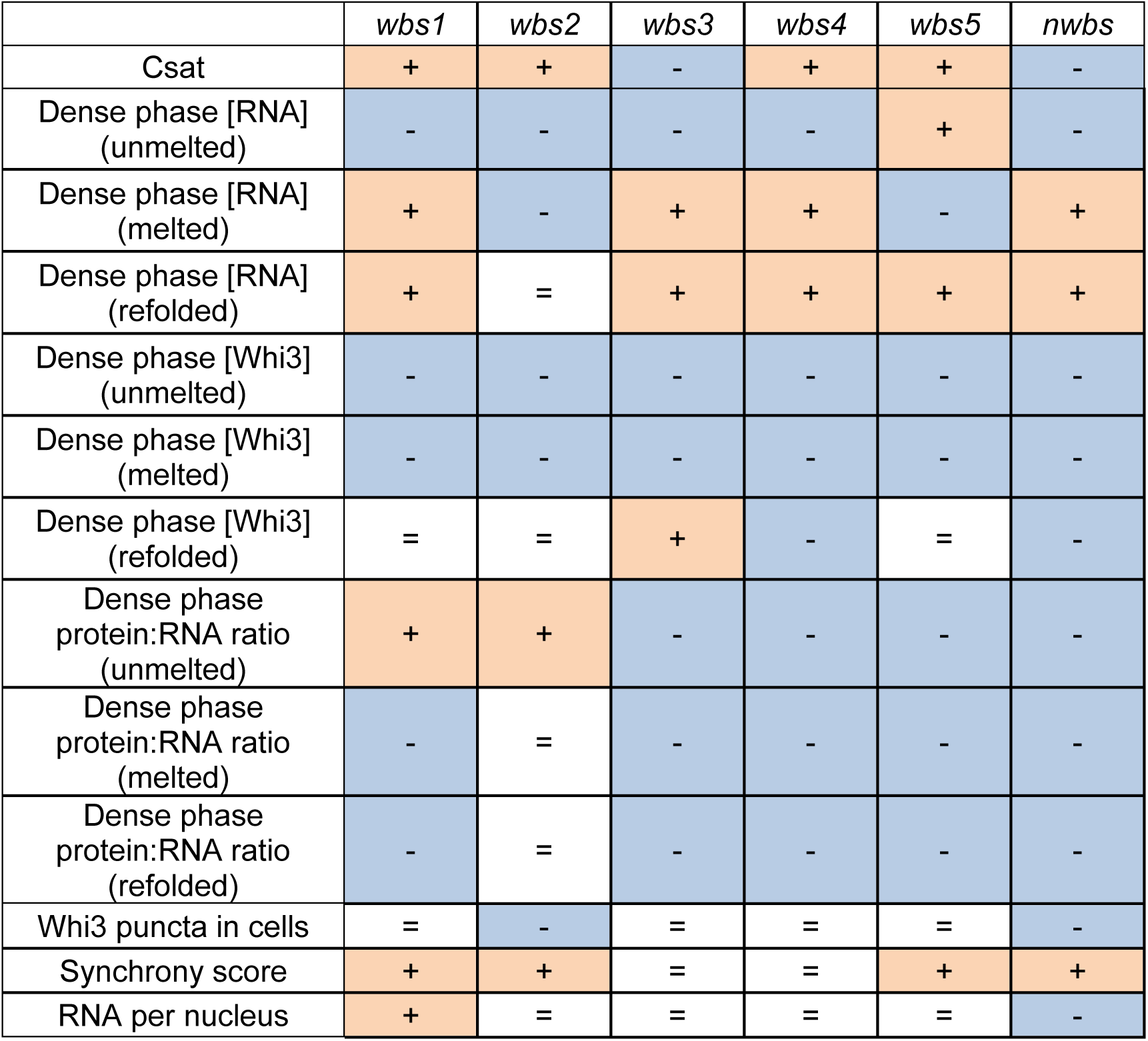
Summary of results comparing mutants to WT with + indicating the value is significantly larger than that of WT, - indicating the value is significantly smaller than WT, and = indicating the value is not significantly different from WT.

Why do individual binding sites vary so greatly in their ability to condense with Whi3 protein? RNA bind-n-seq data from Figure 1 suggest that the surrounding sequence can influence the affinity of Whi3 to a canonical binding site. However, unmutated binding site 1, binding site 3, and binding site 5 share identical immediate flanking nucleotides (having a C directly upstream of the motif and an A directly downstream), yet when mutated do not elicit similar condensate properties. Instead, we see that *wbs3* has a reduced Csat compared to *wbs1* and *wbs5*, and *wbs5* also shows a significant increase in the dense phase RNA concentration. It seems unlikely that other nucleotides beyond those immediately upstream or downstream of the five-nucleotide binding site would contribute to affinity significantly as RRMs typically only recognize a 2-8 nucleotide stretch (Auweter et al., 2006; Cléry et al., 2008). It is possible that *wbs5* behaves differently since it is mutated to UACAC instead of CUAGU, which does have a higher R value, but this still does not explain the difference between *wbs1* and *wbs3*. It is possible that there is cryptic cooperativity in protein recruitment depending on the position of sites relative to one another or other elements of the RNA in 3D that could account for differences not explainable by the immediate context.

The surrounding structure of the RNA may also impact Whi3’s accessibility to binding sites. While the data in Figure 4 did not show a significant change between mutants and WT in RNA structure and ability to dimerize, the average base pairing probability (BPP) for individual binding sites are different from each other which may affect Whi3’s affinity for the site (Figure 4B). This seems to be further supported by the melting data in Figure 5 which shows that melting the RNA to eliminate structure results in the dense phase protein concentration that is more similar to WT than unmelted counterparts. It at first may seem surprising then that the refolded RNA results do not look more like unmelted RNA; however, we believe that this is due to differences in folding pathways since the refolded RNA was folded at 25 °C post-transcription instead of 37 °C during transcription. The overall increased recruitment of components to the dense phase when the RNA is melted suggests that there is also a non-specific role of ssRNA binding that under certain contexts could promote Whi3 binding to more or different target RNAs under stress conditions, consistent with its role in stress granules in *S. cerevisiae* (Cai & Futcher, 2013; Holmes et al., 2013). Together, these data provide strong evidence that RNA folding history and local structure critically shape condensate composition.

Importantly, the mutants do not simply alter *in vitro* condensates, but we can see effects *in vivo* as well. The *wbs1*, *wbs2*, *wbs5* and *nwbs* mutants all show changes from asynchronous nuclear division to more synchronous division. For *wbs2*, this seems to be due to an inability to form condensates since these cells show less Whi3 in puncta yet have similar transcript levels to WT. For the other mutants, we suspect functional impacts are likely due to compositional or material state changes that are difficult to quantify in cells due to the small size of these condensates. Alternatively, the functional impacts could be due to altered Whi3-RNA interactions in dilute-phase complexes. Like *wbs2*, *nwbs* has less Whi3 in puncta, but it also shows lower numbers of RNAs possibly due to overall lower expression levels or increased degradation potentially because of poor Whi3 binding. *wbs1* shows comparable amounts of Whi3 in condensates as WT, yet has more RNA per nucleus which could mean higher expression levels of transcript or that the transcript is less susceptible to degradation than WT. Potentially, this increase in transcript levels could result in more condensates than if the transcript level were the same as WT. Finally, *wbs5* does not show any change in the amount of Whi3 in condensates or in transcript levels, yet it is still more synchronous than WT. This signifies that abrogated *CLN3*-Whi3 interactions can still affect biological function even if it does not eliminate condensates through either altered dense phase properties or a change in an unknown dilute phase function. Finally, there are notable differences between the behavior of mutants *in vivo* and *in vitro* and in some cases (*wbs2*) opposing effects on the propensity to make condensates. These differences are likely due to the increased molecular complexity and regulation *in vivo*.

It was surprising to see that a subset of binding site mutants actually enhanced phase separation at lower protein and RNA concentrations. The *CLN3*-Whi3 condensate system has a balance of both homotypic (protein-protein and RNA-RNA) and heterotypic interactions (protein-RNA). Previous work has shown how changing homotypic interactions within the *CLN3*-Whi3 systems can affect phase behavior. Increasing the interaction propensity of the coiled-coil region in Whi3 does increase oligomerization of Whi3 protein; however, this oligomerization impedes phase separation. In contrast, deleting or weakening the coiled-coil domain enhances phase separation by allowing for more varied and weaker homotypic interactions between the protein (Seim et al., 2022). Similar results have been seen with *CLN3* in which conformation mutants that increase multimerization of RNA result in reduced RNA recruitment to the dense phase and result in a smaller, more branched assemblies (Seim et al., 2024). Modeling experiments show that *CLN3*-Whi3 phase separation is primarily driven by heterotypic interaction, though these interactions can be specific strong interactions or weaker nonspecific interactions (Lin et al., 2023). Evidence of weaker interactions can be seen with the *nwbs* mutant and a mutant version of *BNI1*, another RNA that binds to Whi3, that lack UGCAU motifs but can still form condensates with Whi3 under specific conditions (Lee et al., 2015; Lin et al., 2023). By mutating individual binding sites, potentially stronger or at least more specific interactions between *CLN3* and Whi3 are lost while leaving intact potentially weaker, non-specific interactions. This decrease in the apparent Csat of mutants could be the result of losing specific interactions actually can help promote dilute-phase assemblies or microphases that normally compete with the macroscopic condensates in the wild-type setting. Since RNA bind-n-seq data and structural analysis provides evidence that the binding sites likely have different affinities and Whi3 accessibility, certain sites may contribute more to the stronger, specific contact than others, explaining the differences we see between mutants.

In conclusion, we see that shuffling just 5 nucleotides out of the 1,613 nucleotide sequence of *CLN3* can change condensate formation and disrupt normal cellular function. Through these mutations we are removing a specific interaction occurring between *CLN3* and Whi3 while retaining nonspecific interactions. This balance of interactions strengths has been studied before in the context of homotypic interactions, and it has been shown in these cases that increasing strong interactions between components change condensate properties. This system suggests that similar principles apply to heterotypic interactions. An argument could be made that multivalency in the context of RNA properties is too broad of a term and it is important to elaborate what kind of interactions/contacts are driving phase separation.

## Materials and Methods

### *In vitro* protein prep

Full-length Whi3 was tagged with a 6-His tag at the N terminus and transformed into BL21 bacteria for protein expression using KAN resistance to select for bacteria. The transformants were allowed to grow overnight on one plate before being resuspended with 5 ml of 2XYT media and added to 1 L flasks. Cultures were grown at 37 °C until reaching OD 0.6, then induced with 1 mM isopropyl β-d-1-thiogalactopyranoside and grown for 20 h at 18 °C. Cells were collected by centrifugation and lysed in lysis buffer (1.5 M KCl, 50 mM HEPES, pH 7.4, 20 mM imidazole, 5 mM beta-mercaptoethanol, and one tablet of Roche protease inhibitor cocktail). The lysate was centrifuged to clarify, and the supernatant was incubated with Co-NTA resin. Beads were washed with lysis buffer in a gravity column, and protein was eluted in elution buffer (150 mM KCl, 50 mM HEPES, pH 7.4, 200 mM imidazole, 5 mM beta-mercaptoethanol). Proteins were dialyzed in Whi3 buffer (150 mM KCl, 50 mM HEPES, pH 7.4, and 5 mM beta-mercaptoethanol). Protein purity was confirmed via 4-20% SDS-PAGE gel. Approximately 10% of eluate was taken to be labeled with Atto 488 dye. This was done by incubating protein with 3X molar excess of dye at room temperature for 30 min then dialyzing off remaining dye in Whi3 buffer.

### *In vitro* RNA transcription

*CLN3* RNA was synthesized using the Hi-Scribe T7 kit (New England Biolabs E2040) according to the manufacturer’s instructions. Briefly, 1 μg of DNA template was added to a tube with 1.5 μl of 10x reaction buffer, 1.5 μl of Hi-T7 polymerase, 0.5 μl of RNAse inhibitor, 1.5 μl of each NTP at 100 mM, 0.1 μl of Cy-5 labeled UTP, and enough water to bring the total volume to 20 μl. RNA synthesis was performed on a thermocycler set to incubate at 37 °C for 8 h. Following synthesis, the solution was diluted to 50 μl with RNase-free water, and the DNA was digested by the addition of 2 μl of RNAse-free DNase directly to the tube (New England Biolabs, M0303). RNA was precipitated by the addition of 25 μl of 2.5M LiCl, followed by chilling at -80 °C for 30 min. The RNA was pelleted by centrifugation at maximum speed (20,000rcf) for 10 min, washed briefly with 70% ethanol, and then resuspended in nuclease-free water. The concentration was determined by measuring A260 using a Nanodrop. Purity was assessed by denaturing gel electrophoresis.

### *In vitro* condensate assay

For the in vitro phase separation assays, glass-bottom imaging chambers (Grace Bio-Labs) were blocked with PLL-PEG in Whi3 buffer (150 mM KCL, 50 mM HEPES, 5 mM beta-mercaptoethanol) for 30 min to prevent protein adsorption to the surfaces of the well. The surfaces were washed thoroughly with Whi3 buffer before the addition of 5% Atto488-labeled protein and Cy3-labeled mRNA solutions. Concentrations of protein and RNA were checked via Nanodrop. Whi3 buffer was added, followed by Whi3 protein, followed by RNA to the final concentration indicated in the panel. For melting experiments, prior to RNA being added it was heated at 95 °C for 1 min to melt secondary structure. Refolded RNA was also melted for 1 min at 95 °C then immediately cooled to 25 °C and allowed to refold for 30 min. All wells were mixed and then incubated at 25 °C for 5 h, after which they were imaged at room temperature with a Zeiss LSM 980 with Airyscan 2 using an Apochromat 63x 1.40NA Oil immersion objective. Illumination was done using 488 nm and 639 nm diode lasers. Detection was with a high QE 32-channel spectral array GaAsP. Microscope control and analysis were performed on a premium HP Z6 workstation running Zeiss ZEN 3.7.

### *In vitro* condensate image analysis

To analyze images of condensates, we built a custom Python code that included scikit-image, pandas, numpy, scikit-learn, and scipy (Harris et al., 2020; McKinney, 2011; Pedregosa et al., 2011; Virtanen et al., 2020; Walt et al., 2014). In brief, the code identifies the protein and RNA channel and separates the two. Using the protein channel, a mask was made using Otsu thresholding and water shedding to identify separate objects. The shape, centroid, integrated intensity of the entire object, and the average intensity of pixels including and immediately adjacent to the centroid were measured in both channels. Objects with a volume less than 125 pixels and an average protein centroid intensity less than 1000 were excluded as noise. Each condition was repeated in three different wells, and two images were taken of each well. The 50 largest condensates from each well were used for data analysis. The RNA intensity, protein intensity, and size were all tested for normalcy with the Shapiro-Wilk test. This confirmed that some of data for the different strains did not fit a normal distribution, so differences between groups were determined with the Kruskal-Wallis test and post hoc Dunn Q test.

### FRAP

Condensates were formed as above, but were imaged after 1 h, at which point they are more liquid. Imaging was performed with a Zeiss LSM 980 with Airyscan 2 using an Apochromat 63x 1.40NA Oil immersion objective. The microscope was equipped with an incubation chamber allowing for control of temperature at 25°C. Illumination was done using a 488 nm diode laser set to 0.5% power and maximum scan speed. For FRAP, three images were taken of the condensates before bleaching. Bleaching was performed using the 488 nm laser set to 100% power and a scan speed that was empirically determined to lead to a ∼70% decline in fluorescence. After the bleach step, images were taken every 10 s. Two biological replicates were taken and three wells were used for each biological replicate to act as technical replicates. At least 3 separate condensates were used per well, meaning a total of 18 condensate regions of interest were used to calculate the FRAP curves.

### Cloning and Strain preparation

Mutant *CLN3* constructs were created using classical site directed mutagenesis and transformed into DH5α with plasmids listed in in table below (New England Biolabs). *Ashbya gossypii* integration constructs for WT *CLN3* and *CLN3* with mutations in each of the Whi3 binding sites were constructed using various strategies. First, WT *CLN3* with the endogenous 3’UTR and 3’ homology was constructed. A double stranded DNA fragment was synthesized that contained the *CLN3* 3’ UTR and terminator (Twist Biosciences) that could be used as a PCR template for AGO6386/6387. Then fragments containing a G418 resistance cassette (from AGB1524 using AGO6388/6389) and additional *CLN3* 3’ homology (from Ashbya gossypii genomic DNA with AGO6390/6391) were amplified and Gibson cloned into BstAP1/BamHI digested AGB1524. The resulting plasmid was AGB1603. To make the *wbs1* mutant a PCR fragment was amplified from AGB1258 with AGO6459/6460 and Gibson cloned into with AatII/FspAI digestedAGB1603 resulting in AGB1618. To make the *wbs2* mutant a PCR fragment was amplified from AGB1254 using AGO6622/6623 and Gibson cloned into AatII/BseRI digested AGB1603 resulting in AGB1639.To make *wbs3* and *wbs4* mutant fragments from AGB1608 and AGB1256 were amplified using AGO6463/6464 and cloned into AGB1603 amplified with AGO6461/6462 to make AGB1619 and AGB1620 respectively. To make the *wbs5* mutant PCR fragments were amplified from AGB1251 (using AGO6527/6529) and AGB1603 (using AGO6524/6528) and Gibson cloned resulting in AGB1625. To make the *nwbs* mutant AGB1083 and AGB1625 were digested with AatII/BseRI and ligated using T4 DNA ligase (NEB) resulting in AGB1640.

To make *Ashbya gossypii* strains with eGFP tagged Whi3 fragment from AGB1441 made using BMGB1/Spe1 digestion. The digested fragment was transformed into young mycelia of strain AG416 (ΔlΔt2) by electroporation and plated to AFM + NAT. Resistant mycelia were checked for correct integration using AGO578/500 (for 5’ end) and AGO264/660 (for 3’ end) after which positive fragments were sequenced. Clean spores were individually plated to make homokaryons and recorded as AGB977. *Ashbya gossypii* strains with *CLN3* mutants were made using a PCR fragment containing the *CLN3* mutant construct was amplified using AGO2049/3536 for each of the *wbs* mutant plasmids. The PCR fragment was transformed into young mycelia of strain AG977 (WHI3-eGFP NAT) by electroporation and plated to AFM + G418 (G418; GoldBio) (Altmann-Jöhl & Philippsen, 1996). G418 resistant mycelia were checked for correct integration by PCR amplification using AGO432/6393 (for 5’ end) and AGO6304/6394 (for 3’ end). PCR positive 5’ fragments were additionally sequenced to verify that the desired *wbs* mutation was present in the clone. *CLN3* mutant strains were grown from heterokaryon spores under selection to ensure expression of mutant gene.

Primers:

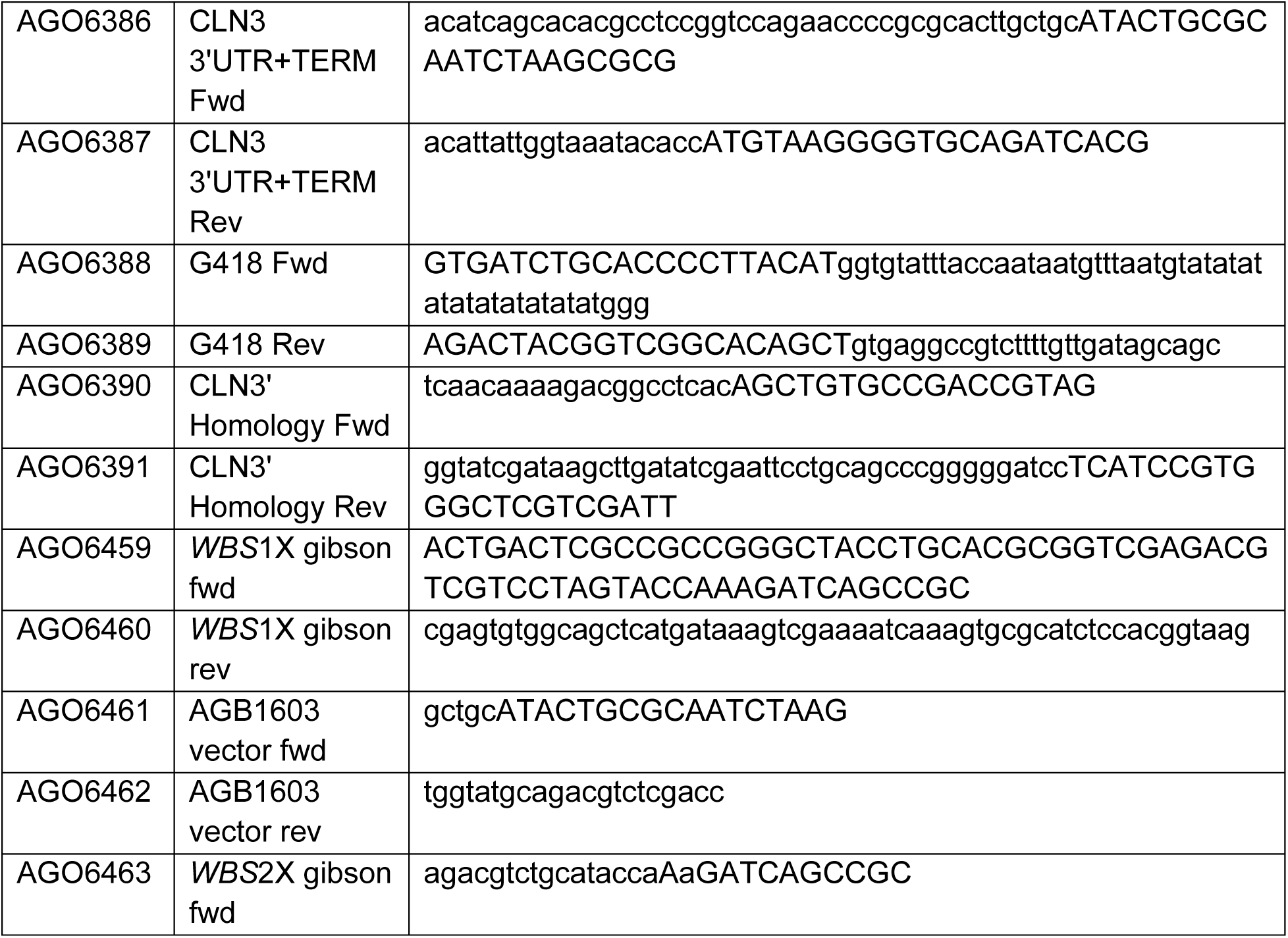
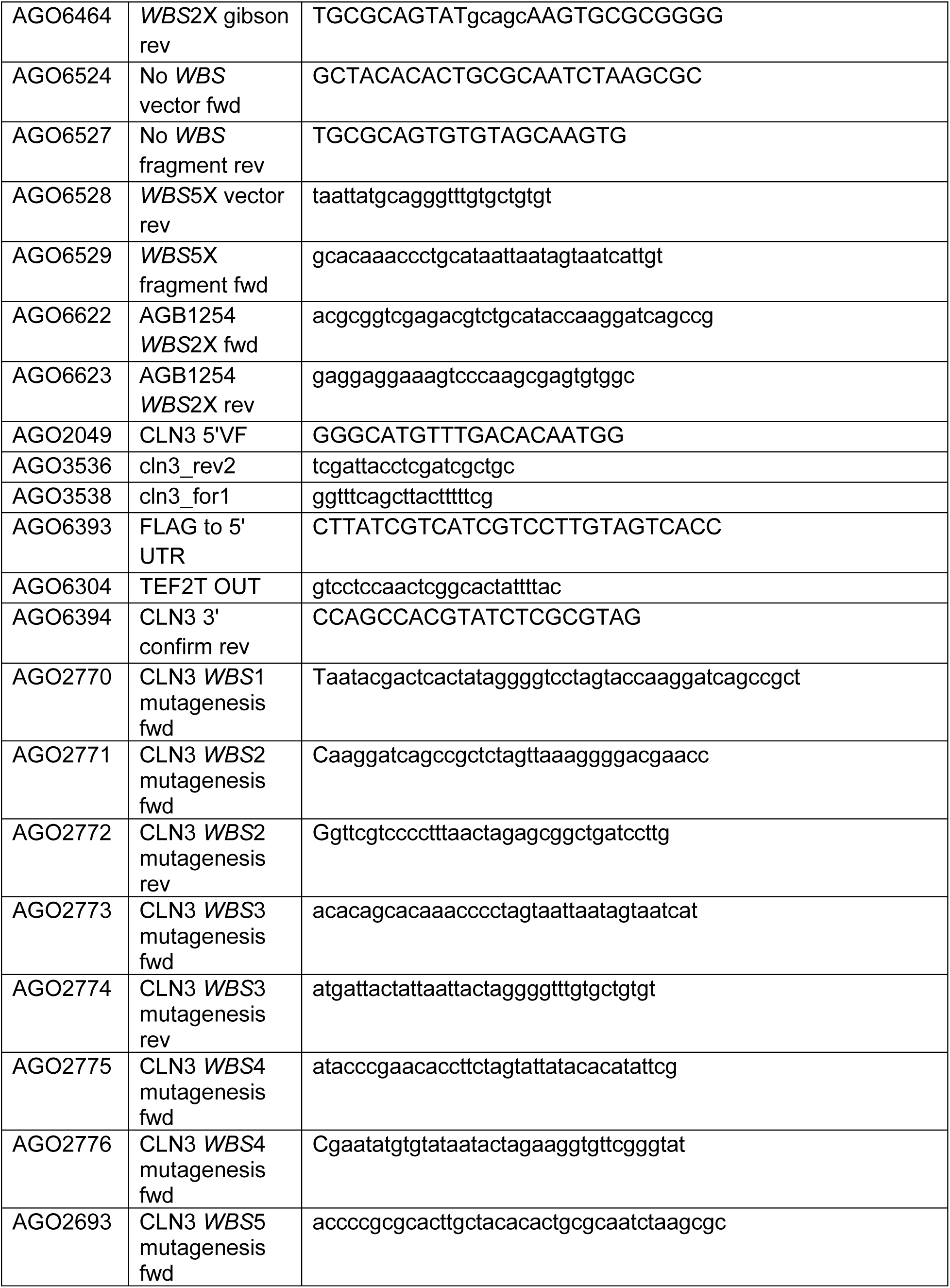
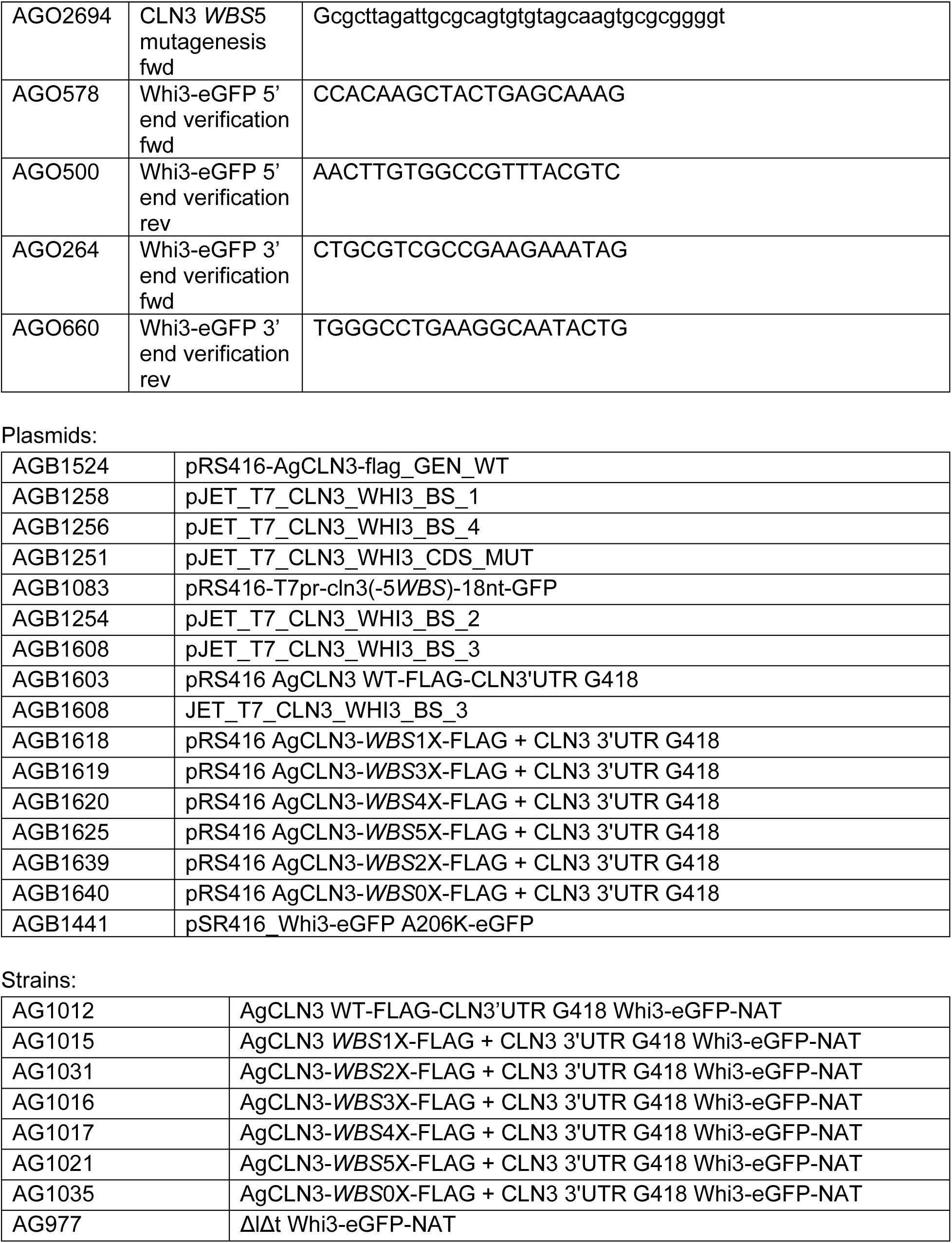

### RNA-FISH and IF

Single-molecule RNA FISH was conducted based on previously published protocols (Lee et al., 2013, 2016). Cells were grown for 16 h at 30 °C in liquid *Ashbya* Full Media (AFM, 10 g Tryptone, 10 g Yeast Extract, 1 g Myo-inositol, 50 ml 40% Dextrose, in 1 L Water), then fixed with formaldehyde at a final concentration of 3.7% vol/for 1 h. Mycelia were then gently pelleted by centrifugation (400xcf) and washed twice with ice-cold buffer B (1.2 M sorbitol and 0.1 M potassium phosphate, pH 7.5). The fixed cells were resuspended in 1 ml spheroplasting buffer (10 ml buffer B and 2 mM vanadyl ribonucleoside complex). To remove the cell wall and permeabilize the cells 1.5 mg/ml Zymolase, prepared with RNAse-free water, was added, and the mycelia were incubated at 37 °C for around 20 min. Individual digestion time varies, so digestion was monitored by DIC microscopy every 5 min; permeabilization was considered complete when the cells became phase dark. After digestion, cells were immediately washed twice with buffer B, then resuspended in 1 ml RNase-free 70% ethanol for 4 h at 4 °C.

Custom TAMRA labelled RNA FISH probes (Biosearch Technologies) were designed and were resuspended in 20 μl 1x TE buffer. Following the 4 h incubation in ethanol cells were washed with wash buffer (2x SSC (0.3 M Sodium Chloride, 30 mM trisodium Citrate), 10% vol/vol deionized formamide), resuspended in 100 μl hybridization buffer (100 mg/ml dextran sulfate, 1 mg/ml E. coli tRNA, 2 mM vanadyl ribonucleoside complex, 2 mg BSA, 2x SSC, and 10% vol/vol deionized formamide in 10 mL final volume with RNAse-free water) with 25 nM mRNA FISH probe and 1:200 Rat anti Tub antibody, then incubated in the dark, overnight, at 37 °C. On the second day, cells were washed 3 times with 1 ml of 1 mg/ml BSA in RNase-free PBS. After the third wash, resuspended cells in 100 μL of 1 mg/ml BSA in RNase-free PBS with 1:500 Hoechst and 1:500 goat anti-rat 488. Allowed this to incubate at room temperature for 1 h out of the light. After the hour incubation, the cells were washed 3 times again with 1ml of 1 mg/ml BSA in RNase-free PBS without any stains or antibodies. Cells were mounted with Vectashield. covered with an RNase-free coverslip, sealed with nail polish, and imaged using a Nikon Ti-E stand equipped with a Yokogawa CSU-W1 spinning disk confocal unit, a Plan-Apochromat 100x/1.49 NA oil-immersion objective, and a Zyla sCMOS camera (Andor). Nikon NIS-Elements software v.4.60 was used to control the machine. The spinning disk was illuminated with 405 nm, 488 nm, and 561 nm laser source. This was done on two independent transformants for each strain.

### Synchrony Scoring

Images were scored by determining the cell cycle stage for each nucleus, where possible, based on the tubulin IF staining pattern. In *Ashbya,* the spindle pole body forms a single focus in G1, a pair of juxtaposed foci in S/G2, and two foci connected by a spindle in M-phase (Rüthnick & Schiebel, 2016). We counted the total number of nuclei in each cell cycle stage. We also counted all pairs of adjacent nuclei and counted the number of adjacent nuclei that were at the same stage of the cell cycle. We then calculated a synchrony score, which was the actual number of neighboring nuclei in the same cell cycle state divided by the number of such pairs expected by chance. If the nuclei behave independently of each other, this number will equal 1, with higher numbers indicating nuclei are more synchronous with their neighbors than expected by chance. A minimum of 200 nuclei were scored for each transformant.

### RNA-FISH analysis

Individual images were cropped, z-projected, and contrasted manually in ImageJ. These cropped images were used to train a model in iLastik (Berg et al., 2019). Briefly, an iLastik model was trained on a subset of the data (∼2% of the total data) to recognize nuclei and FISH foci. Following training, the model was used to segment all the images in the set. The segmentation was manually checked for each image, and those that failed this QC check, due to an abundance of false positives or negatives, were removed from the data set. Following segmentation, the segmented files were used to map Regions of Interest (ROIs) onto the original files. These regions of interest were used to measure the position, size, and fluorescence intensity of the nuclei and FISH foci. To compare amounts of FISH foci in cells the ratio of foci: nuclei was taken. This data was tested for normalcy with the Shapiro-Wilk test. This confirmed that the data for the different strains did not fit a normal distribution, so differences between groups were determined with the Kruskal-Wallis test and post hoc Dunn Q test.

### Live cell imaging

*Ashbya* strains were grown in 25 mL of liquid AFM for 16 h under double G418 and Nat selection. 10 mL of culture was spun down at 400 rpm for 5 min, and the media was removed. The pellet was washed twice with a solution of 50 mM Tris and 150 mM NaCl. After the second wash, approximately 20 μl of the suspension was placed on a slide, and a coverslip was placed on top, which was sealed with VALAP. Cells were imaged within 30 minutes of mounting using a Nikon Ti-E stand equipped with a Yokogawa CSU-W1 spinning disk confocal unit, a Plan-Apochromat 100x/1.49 NA oil-immersion objective, and a Zyla sCMOS camera (Andor). Cells were illuminated with a 488 nM laser. For image analysis, hyphae were segmented by hand in ImageJ. These images were run through a macro that used a 3D object counter to identify the cell and foci. To calculate the percent Whi3 signal in foci, the sum intensity of the foci was divided by the sum intensity of the total Whi3 signal in the cell. This data was tested for normalcy with the Shapiro-Wilk. This confirmed that the data for the different strains did not fit a normal distribution, so differences between groups were determined with the Kruskal-Wallis test and post hoc Dunn Q test.

### Four-base DMS-MaP

Structure probing was done similar to previously published methods using conditions suitable for DMS modification of all four nucleotides (Dey et al., 2024; Mitchell et al., 2023). Briefly, *in vitro*–transcribed RNAs were prepared as previously described and diluted to 5 µg total RNA in 25 µL nuclease-free water. RNA was denatured at 98 °C for 1 min and snap-cooled on ice for 1–2 min. Folding was initiated by the addition of 25 µL of 2× folding buffer (600 mM bicine pH 8.3, 200 mM NaCl, 10 mM MgCl_2_), yielding final concentrations of 300 mM bicine, 100 mM NaCl, and 5 mM free Mg^2+^. Samples were incubated at 37 °C for 30 min before chemical probing. DMS was diluted 1:5 into ethanol to generate a 1.7 M working solution. Working solutions (1 µL) of DMS and ethanol control were aliquoted into individual tubes and rapidly mixed with 9 µL of Folded RNA. Reactions were incubated at 37 °C for 3 min for ethanol controls and 6 min for DMS. DMS reactions were quenched by the addition of 10 µL of freshly prepared 25% β-mercaptoethanol. All reactions were immediately placed on ice. Probed RNAs were buffer exchanged using MicroSpin™ G-50 spin columns (Cytiva, USA).

Reverse transcription was performed under mutational profiling conditions optimized for DMS probing (Smola et al., 2015). Purified RNA (8.8 µL) was combined with random nonamer RT primers (NEB S1254, 200 ng/uL) and dNTPs, heated to 98 °C for 1 min, and snap-cooled on ice. Reverse transcription reaction buffer was added, with the following final concentrations: 1× NTP-minus first-strand buffer (0.5 M Tris pH 8.0, 0.75 M KCl, 0.1 M DTT), 1 M betaine, and 6 mM MnCl_2_. SuperScript II reverse transcriptase was added to experimental samples, and reactions were incubated at 25 °C for 10 min, followed by 90 min at 42 °C with periodic temperature cycling, and enzyme inactivation at 70 °C for 10 min. First-strand cDNA was purified using G-50 spin columns. Double-stranded cDNA was generated using the NEBNext Ultra II Non-Directional RNA Second Strand Synthesis Module (NEB E6111S). Entire first-strand reactions were used as input and incubated for 3 h at 16 °C. Reactions were purified using Agencourt AMPure XP beads at a 1.8× bead-to-sample ratio and eluted in nuclease-free water. DNA concentration was quantified using the Qubit dsDNA High Sensitivity assay. Purified cDNA was converted into sequencing libraries using the NEBNext Ultra II FS DNA Library Prep Kit for Illumina. Fragmentation, end repair, and adapter ligation were performed according to manufacturer guidelines, with fragmentation time optimized for each RNA to achieve desired insert sizes. Libraries were PCR-amplified using indexed primers and Q5 Master Mix, followed by bead-based cleanup. Final libraries were quantified by Qubit and assessed for size distribution using an Aligent TapeStation. The libraries were sequenced on the Illumina NextSeq1000.

### DMS-MaP analysis

Chemical probing data were analyzed using ShapeMapper2.2.0 with DMS mode (-dms) (Busan & Weeks, 2018). ShapeMapper was run to parse and count mutation events from sequencing reads, producing both per-nucleotide mutation rates and per-read histograms. Parsed and counted mutation files were retained for downstream analysis. Default quality filtering parameters were used unless otherwise specified. Structures were folded using population-averaged DMS reactivities with a maximum base-pairing distance of 600 nucleotides. Processed DMS reactivity profiles were visualized using RNAvigate (Irving & Weeks, 2024).

### RNA bind-n-seq with CLN3 oligos

An oligonucleotide pool was amplified by PCR using primers MC1780.5p.RBNS (5′-TAATACGACTCACTATAGGGAGTTCTACAGTCCGACGATC-3′) and MC1781.3p.RBNS.RT (5′-GCCTTGGCACCCGAGAATTCCA-3′). Reactions were performed using Kapa HiFi DNA Polymerase with GC Buffer (Kapa KK2502; Roche 07958897001) according to kit directions. Products were gel-purified and eluted in 40 µl elution buffer. DNA concentration was measured using a Qubit dsDNA Broad Range assay, and samples with ≥20 ng/µl were used for downstream applications.

RNA was generated using the T7 RiboMAX™ Express Large Scale RNA Production System (Promega P1320). Two 100 µL reactions were prepared, each containing 50 µl of 2× T7 buffer, purified PCR product, nuclease-free water to volume, and 20 µl T7 enzyme mix. Reactions were incubated at 37 °C for 4 h, followed by DNase treatment with 1 µl RQ1 DNase per 20 µl reaction for 15 min at 37 °C. Reactions were pooled, extracted with acidic phenol–chloroform, followed by chloroform–isoamyl alcohol extraction, and RNA was precipitated using sodium acetate, linear acrylamide, and isopropanol. Pellets were washed with 80% ethanol, air-dried, and resuspended in nuclease-free water. The RNA was purified by 8% TBE–urea PAGE. RNA bands were visualized using SYBR Gold staining, excised, and crushed through a pierced 0.5 mL tube by centrifugation. RNA was eluted overnight at 4 °C in gel extraction buffer (20 mM Tris-HCl pH 7, 250 mM NaCl) supplemented with RNase inhibitor. Eluate was filtered through NanoSep columns, and RNA was recovered by phenol–chloroform extraction and ethanol precipitation.

Binding reactions were performed in RBNS binding buffer (25 mM Tris pH 7.5, 150 mM KCl, 3 mM MgCl_2_, 0.01% Tween-20, 500 µg/mL BSA). His-tag Dynabeads were washed three times and equilibrated with recombinant protein for 30 min at 4 °C. RNA was added to a final concentration of 1 µM, with protein concentrations ranging from 10 to 200 nM. Reactions were incubated for 1 h at 4 °C with intermittent mixing. Complexes were washed three times in wash buffer and eluted using 250 mM imidazole in PBS. RNA was purified by phenol–chloroform extraction and ethanol precipitation recovered from RBNS, and input samples were reverse transcribed using SuperScript III (Thermo Fisher 18080093) with primer MC1781.3p.RBNS.RT. cDNA synthesis was performed at 55 °C for 1 h and terminated at 70 °C for 15 min. cDNA abundance was assessed by qPCR using Power SYBR Green and primer MC1782.RP1 with indexed primers. RBNS libraries were amplified using Kapa HiFi HotStart polymerase with Illumina RNA PCR index primers. PCR cycle number (10–16 cycles) was empirically determined based on qPCR results. Libraries were purified, quantified by Qubit and used for downstream sequencing using Illumina NextSeq.

Raw RBNS sequencing reads were processed using a high-performance computing cluster running a Linux operating system. All analyses were performed in Conda-managed environments to ensure reproducibility. Trimmed reads were aligned to the reference RNA sequence using a custom STAR genome index generated from the full-length target RNA sequence (including untranslated regions) (Dobin et al., 2013). Alignments were performed using STAR default alignment parameters. Read counts corresponding to the reference RNA were extracted from STAR alignment outputs for downstream analysis.

### RNA bind-n-seq with randomized oligos

Template DNA was synthesized by IDT and was *in vitro* transcribed using T7 RiboMAX Express large scale RNA kit (Promega) according to manufacturer protocols. The binding, washing, and sequencing protocol was kept the same as the one listed above. R values for 5- and 6-mers were calculated as the frequency of each 5- or 6-mer in the protein-bound sample divided by the frequency in the input pool. Sequence logos were generated by selecting the top 25 5- or 6-mers in R (v. 4.1.0) with R package “ggseqlogo” (Wagih, 2017). RNA secondary structure was predicted with RNAfold based on the random RNA sequences with adaptor sequences. Specifically, a 5- or 6-nt sliding window was generated along each sequence to calculate the average base-pairing probabilities (BPP) for each 5- or 6-mer (Gruber et al., 2008; Lorenz et al., 2011). The average BPP values were then normalized to the corresponding values in the input pool. The normalized BPP values indicate the structural binding bias of each 5- or 6-mer.

## Supporting information

Supplemental Figure 2. Mutants show unique differences in condensate properties in vitro

Supplemental Figure 1. Determining primary RNA interaction sites for Whi3.

Supplemental Figure 3. Sample images of condensates with melted RNA and refolded RNA.

Supplemental figure 3. Normalized DMS reactivity profile of all CLN3 strains with binding sites highlighted with grey boxes, start codon highlighted i

## Acknowledgments

We would like to thank the broader RNA community of the research triangle for supporting collaboration and discussions of this work along with helpful feedback from the Gladfelter lab. This work was supported by Air Force Office of Scientific Research (grant FA9550-20-1-0241) to A.S.G., U.S. National Institutes of Health grants 7R01GM081506-13 and R35 GM156800 to A.S.G, U.S. National Institutes of Health grants R01 HL111527, R35 GM140844 (to A.L.).

## Supplemental Figures and Tables

**Supplemental Figure 1.**
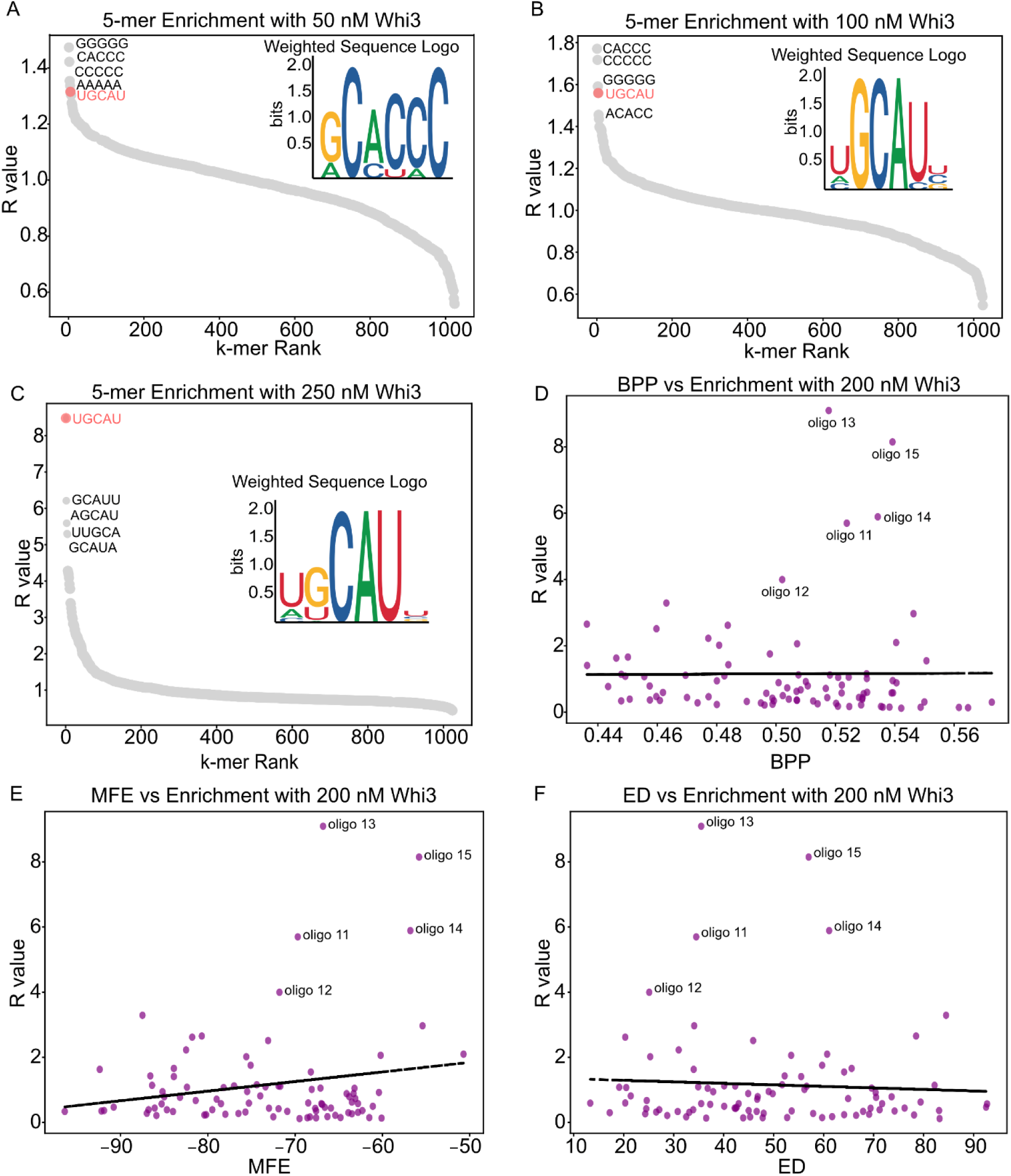
Determining primary RNA interaction sites for Whi3. A) A graph showing which 5 nucleotide motifs were most enriched in the random oligo pool RBNS experiment at 50 nM Whi3, with UGCAU containing motifs highlighted in red. B) A graph showing which 5 nucleotide motifs were most enriched in the random oligo pool RBNS experiment at 100 nM Whi3. C) A graph showing which 5 nucleotide motifs were most enriched in the random oligo pool RBNS experiment at 250 nM Whi3. D) Pearson correlation of *CLN3* oligo base pair probability and R value score. Pearson’s r = 0.01 p-value = 0.954 E) Pearson correlation of *CLN3* oligo minimum free energy (MFE) predictions and R value score (Gruber et al., 2008). Pearson’s r = 0.19 p-value = 0.0764 F) Pearson correlation of *CLN3* oligo ensemble diversity (ED) predictions and R value score (Gruber et al., 2008). Pearson’s r = -0.06 p-value = 0.588

**Supplemental Figure 2.**
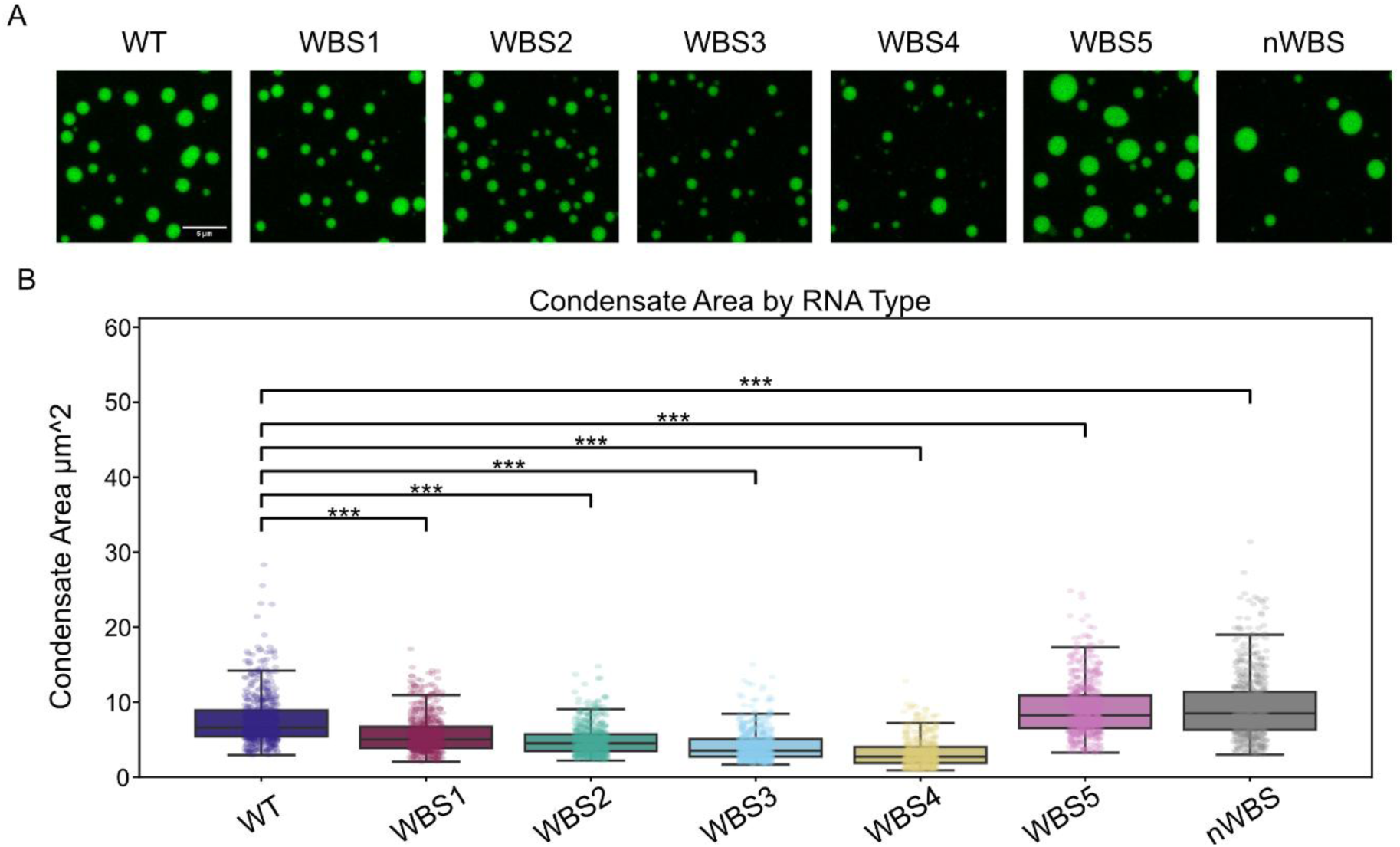
Mutants show unique differences in condensate properties *in vitro* A) Sample images of *in vitro* condensates with different RNA constructs B) Area of max projection images of condensates. Significance calculated using the Kruskal-Wallis test and Dunn Q test *** p < 0.001.

**Supplemental figure 3.**
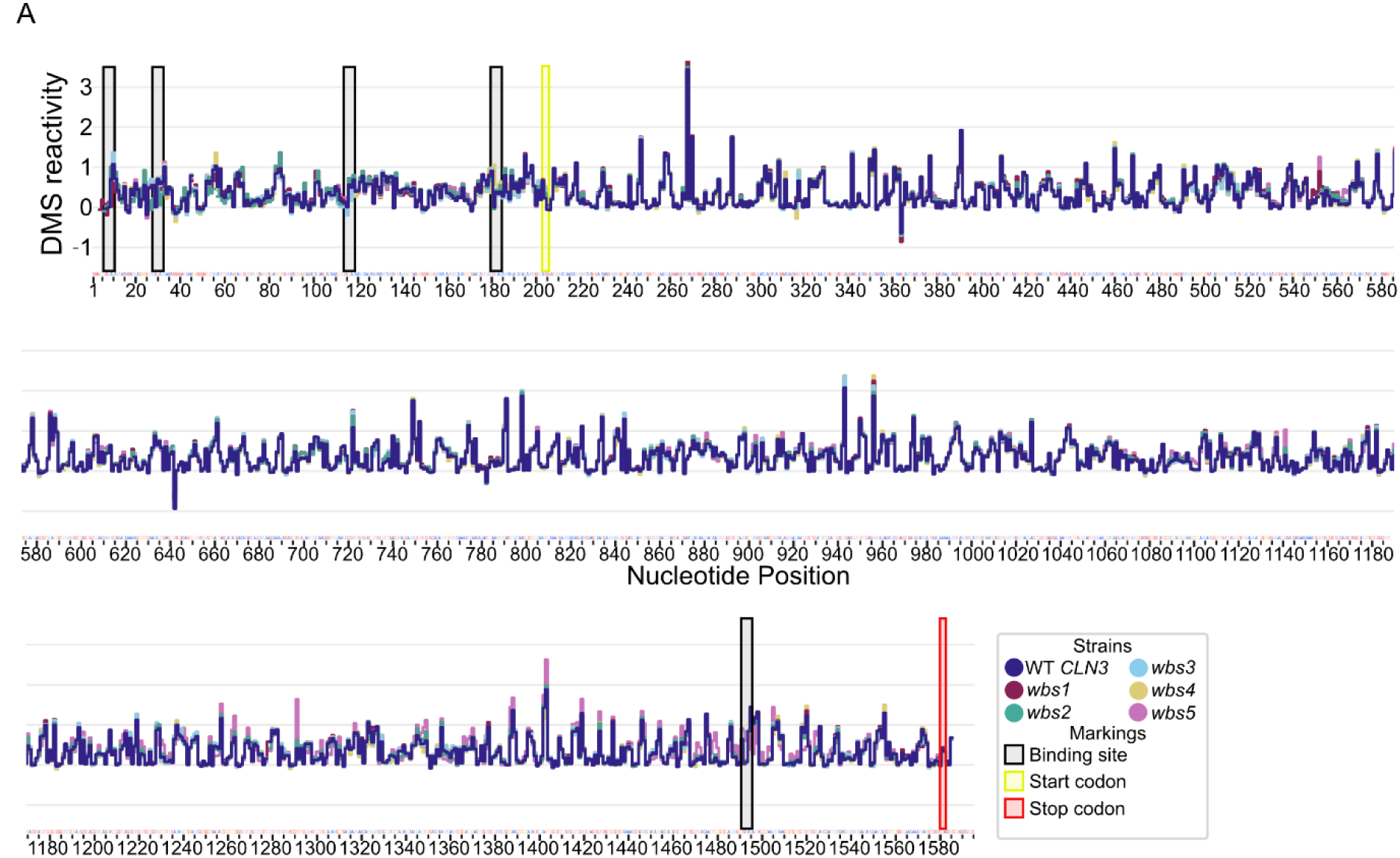
Normalized DMS reactivity profile of all *CLN3* strains with binding sites highlighted with grey boxes, start codon highlighted in yellow box, and stop codon highlighted in red box.

**Supplemental Table 1:**
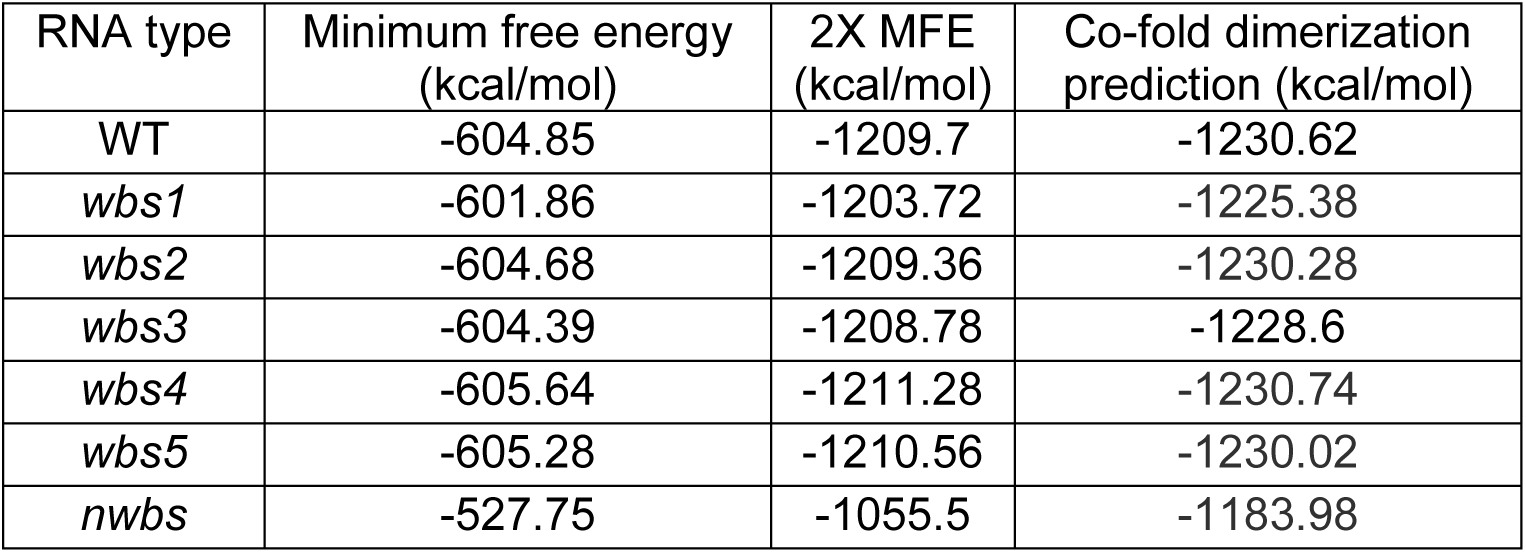
Minimum free energy values predicted using Vienna RNA and predicted co-fold dimer minimum free energy at 25 °C and 150 mM salt concentration (Gruber et al., 2008).

| RNA type | Minimum free energy (kcal/mol) | 2X MFE (kcal/mol) | Co-fold dimerization prediction (kcal/mol) |
| --- | --- | --- | --- |
| WT | -604.85 | -1209.7 | -1230.62 |
| *wbs1* | -601.86 | -1203.72 | -1225.38 |
| *wbs2* | -604.68 | -1209.36 | -1230.28 |
| *wbs3* | -604.39 | -1208.78 | -1228.6 |
| *wbs4* | -605.64 | -1211.28 | -1230.74 |
| *wbs5* | -605.28 | -1210.56 | -1230.02 |
| *nwbs* | -527.75 | -1055.5 | -1183.98 |

**Supplemental Figure 4.**
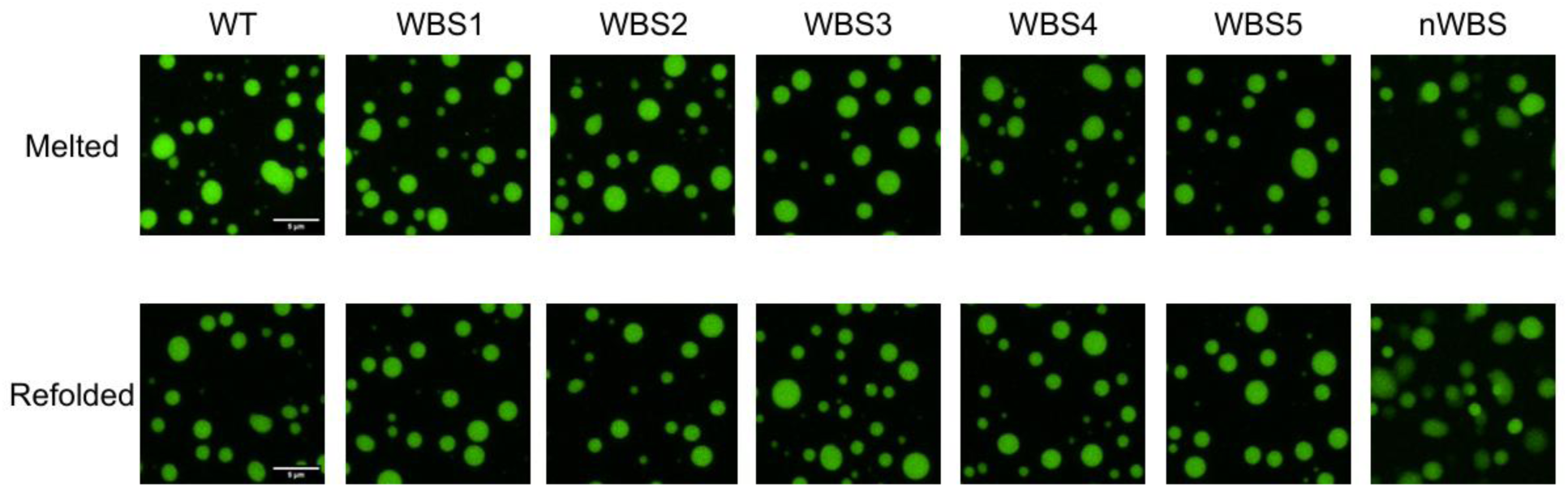
Sample images of condensates with melted RNA and refolded RNA.

