## Supplementary figures and images for "Sequence and structure of protein binding sites in RNA impact biomolecular condensates"

### Supplemental Figure 1. Determining primary RNA interaction sites for Whi3.

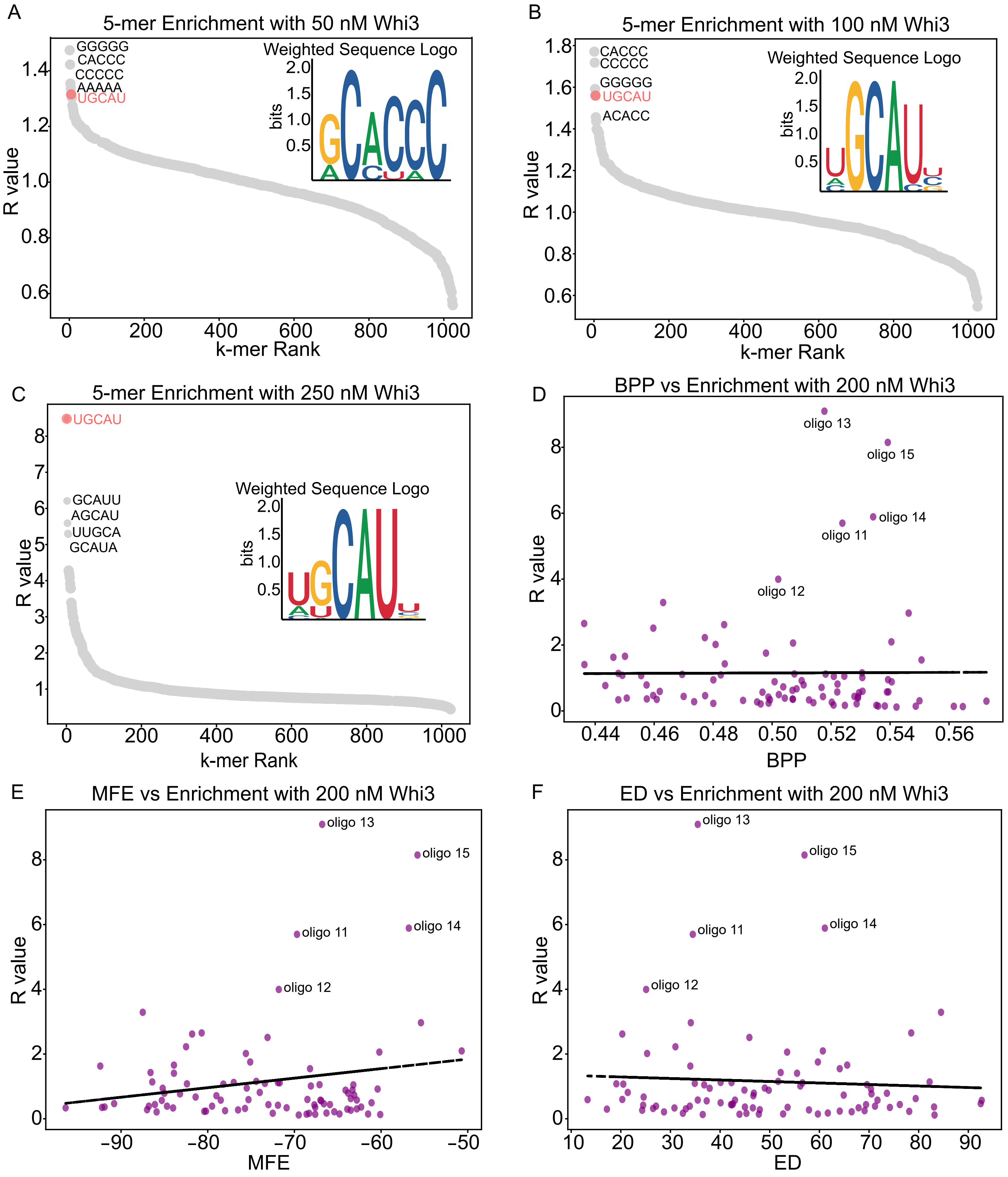

### Supplemental Figure 2. Mutants show unique differences in condensate properties in vitro

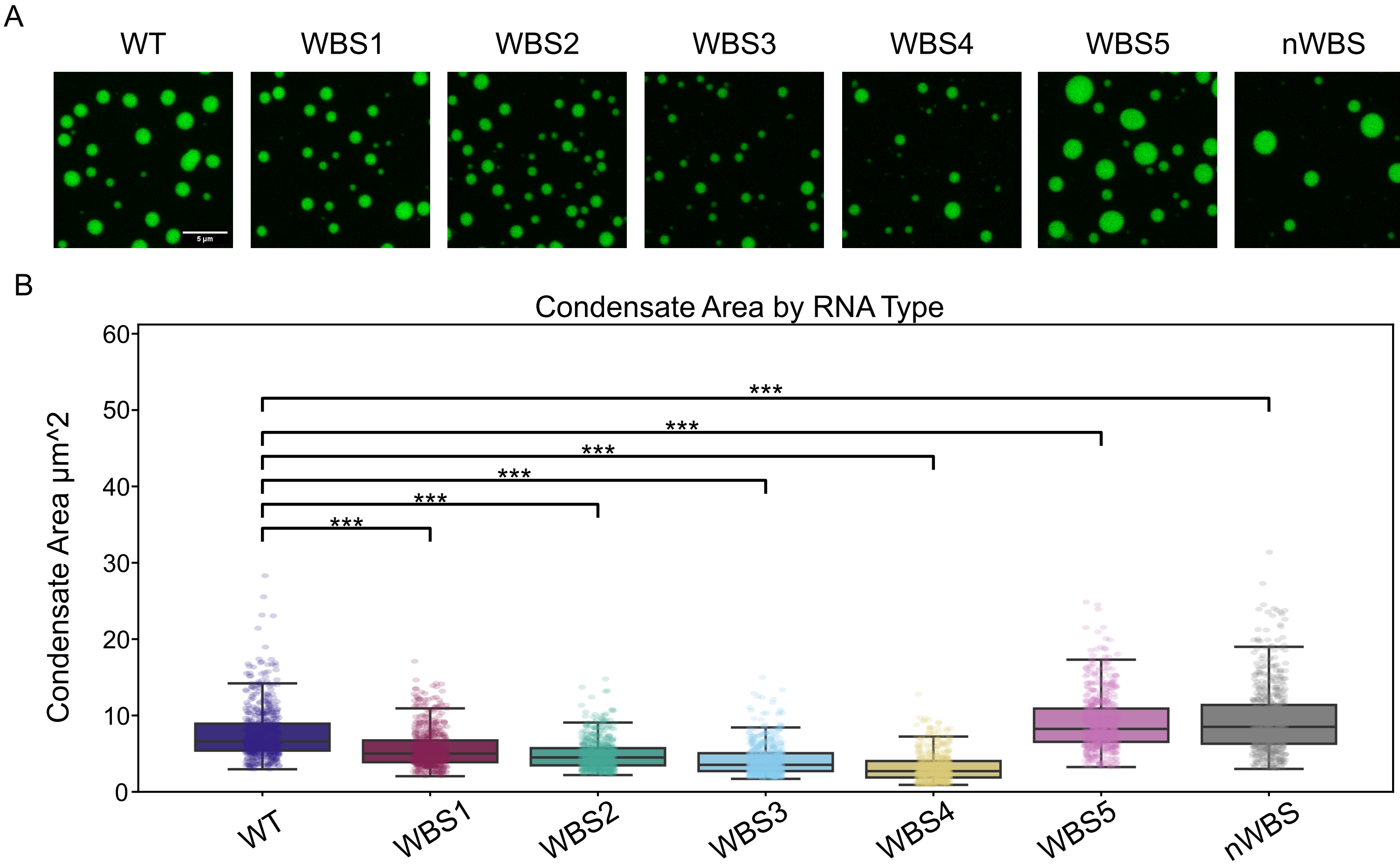

### Supplemental figure 3. Normalized DMS reactivity profile of all CLN3 strains with binding sites highlighted with grey boxes, start codon highlighted i

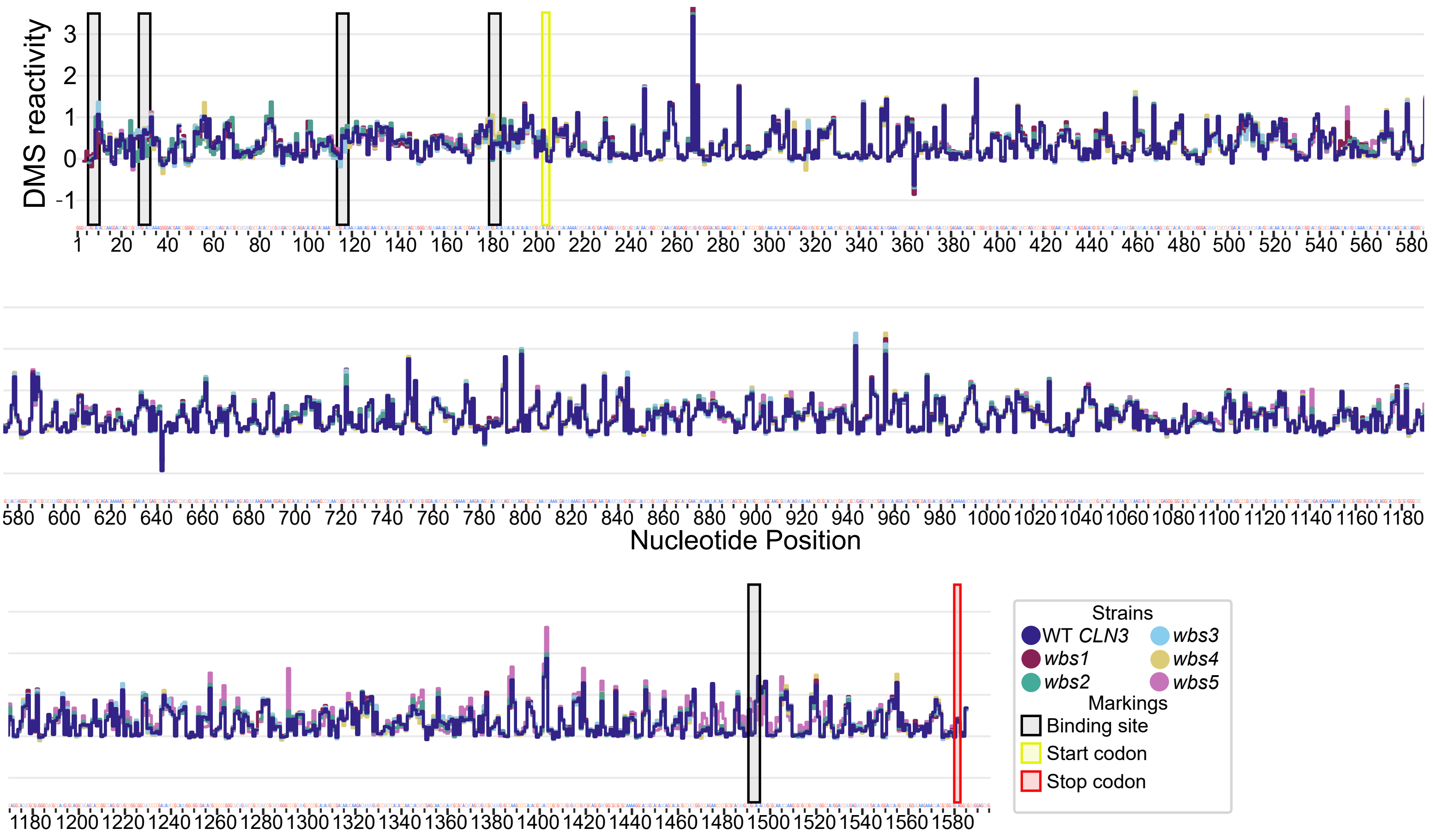

### Supplemental Figure 3. Sample images of condensates with melted RNA and refolded RNA.

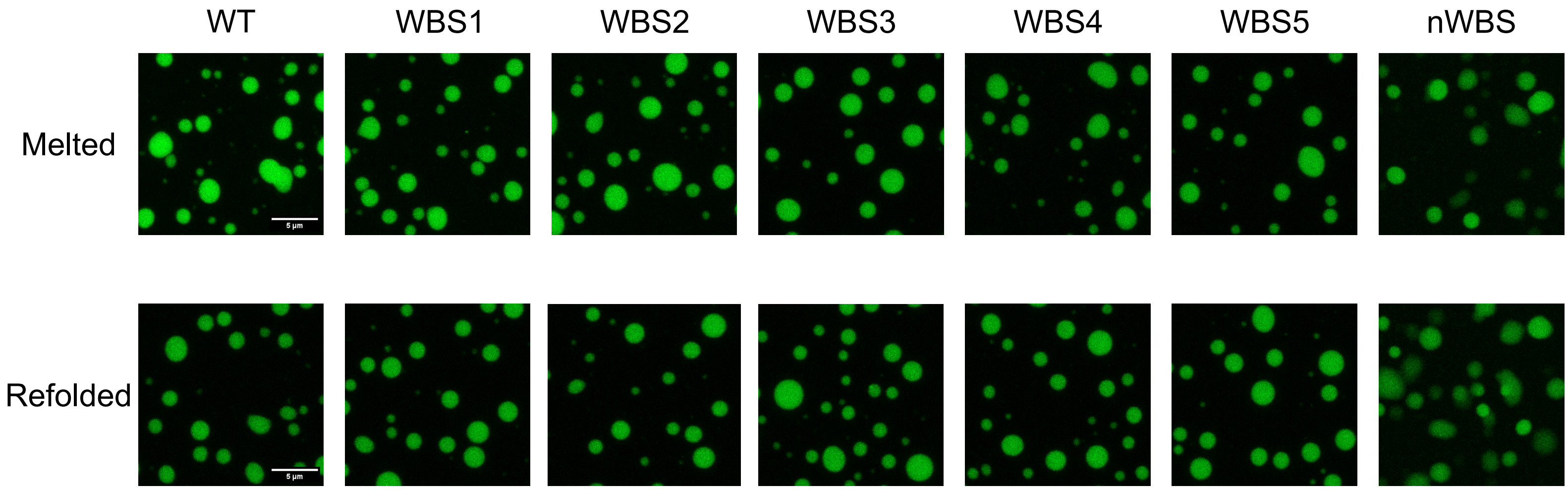
